# Transcription factors recognize DNA shape without nucleotide recognition

**DOI:** 10.1101/143677

**Authors:** Md. Abul Hassan Samee, Benoit G. Bruneau, Katherine S. Pollard

## Abstract

We hypothesized that transcription factors (TFs) recognize DNA shape without nucleotide sequence recognition. Motivating an independent role for shape, many TF binding sites lack a sequence-motif, DNA shape adds specificity to sequence-motifs, and different sequences can encode similar shapes. We therefore asked if binding sites of a TF are enriched for specific patterns of DNA shape-features, *e.g.,* helical twist. We developed ShapeMF, which discovers these shape-motifs *de novo* without taking sequence information into account. We find that most TFs assayed in ENCODE have shape-motifs and bind regulatory regions recognizing shape-motifs in the absence of sequence-motifs. When shape- and sequence-recognition co-occur, the two types of motifs can be overlapping, flanking, or separated by consistent spacing. Shape-motifs are prevalent in regions co-bound by multiple TFs. Finally, TFs with identical sequence motifs have different shape-motifs, explaining their binding at distinct locations. These results establish shape-motifs as drivers of TF-DNA recognition complementary to sequence-motifs.

## Introduction

Diverse cellular processes, including gene regulation, chromatin organization, provirus activity, and DNA replication, depend upon proteins binding to specific genome sites, either alone or in complexes with other molecules. Protein-DNA recognition is thus fundamentally important and critically informs studies of development, disease, and evolution. Protein bound regions can be measured in living cells via imaging and genomic techniques, such as chromatin immunoprecipitation followed by sequencing (ChIP-Seq). To pinpoint binding sites within bound regions, predict binding in the absence of experimental measurements, and shed light on binding specificity, a variety of *in vitro* binding affinity assays and sequence analyses have been deployed.

The specificity of protein-DNA recognition is commonly approached as a problem of discriminating bound nucleotide sequences from other sequences (Bailey 2011, Arvey, Agius et al. 2012, Ghandi, Lee et al. 2014, Setty and Leslie 2015). While most methods focus on enriched patterns of nucleotides, the resulting sequence-motifs become more predictive of *in vitro* and *in vivo* binding when supplemented with structural data based on shape features of the bound DNA (Zhou, Shen et al. 2015, Mathelier, Xin et al. 2016). However, the role of DNA shape in binding specificity has not been investigated outside the context of sequence-motifs. Supporting the hypothesis that DNA-binding proteins can recognize DNA structure without nucleotide recognition, transcription factors (TFs) can bind sequences that do not match sequence-motifs identified using sequence-based searches (von Hippel, Revzin et al. 1974, von Hippel and Berg 1986, Berg and von Hippel 1987, von Hippel 2007, Wang, Zhuang et al. 2012, Yip, Cheng et al. 2012, Afek, Schipper et al. 2014, Slattery, Zhou et al. 2014). Furthermore, the best sequence-based discriminative methods fail to identify a subset of regions bound by TFs, including validated and predicted regulatory elements and regions harboring polymorphisms that correlate with gene expression. Finally, different nucleotides can encode similar DNA structure, so shape features have the potential to be complementary to nucleotide features (Garvie and Wolberger 2001). We therefore sought to investigate the independent role of DNA shape in protein-DNA binding.

To explore how frequently proteins recognize DNA structure and test the idea that structural specificity can occur in the absence of nucleotide recognition, we pose a novel question: are the binding sites of a protein enriched for characteristic patterns of DNA shape features, such as roll, helical twist, propeller twist, and minor groove width? If so these “shape-motifs” might explain many aspects of DNA recognition that are not accounted for by sequence-motifs alone or by DNA shape profiles within sequence-motifs. These open problems include binding to regions that lack sequence-motifs, differential binding of proteins that have very similar sequence-motifs, and low sequence information content positions in and flanking sequence-motifs. To investigate these questions, we developed a novel Gibbs sampling algorithm to discover shape-motifs *de novo* without conditioning on the presence of a sequence-motif. Applying this method to more than 100 TFs and several different cell types, we find that most TFs recognize DNA shape independent of nucleotide recognition.

## Results

### Many strongly bound regions lack a sequence motif

To motivate the need for shape-motifs, we first quantified the prevalence of TF binding without nucleotide recognition. We called sequence motif hits for 110 TFs from regions they bind in human ENCODE data from K562 (chronic myelogenous leukemia) and Gm12878 (lymphoblastoid) cell lines. To broadly define sequence motifs for each TF, we used a collection of position weight matrices (PWMs) from JASPAR (Sandelin, Alkema et al. 2004), TRANSFAC (Matys, Fricke et al. 2003), *in vitro* studies (Berger, Philippakis et al. 2006, Berger, Badis et al. 2008, Badis, Berger et al. 2009, Jolma, Yan et al. 2013), and *de novo* motif-discovery methods, as were curated in (Kheradpour and Kellis 2014), plus up to five *de novo* motifs per TF that we learned from its top 2000 peaks using gkmSVM (Ghandi, Lee et al. 2014). Combining hits to any of these PWMs, we found that a large fraction of the top 2000 peaks for each TF lack a sequence motif for that TF (Table S1). For a typical TF, 29.6 of peaks have no sequence motif. This fraction varies significantly across TFs (range = 0.4-98), but is fairly consistent between cell lines. These findings show that factors other than direct nucleotide recognition are likely influencing TF DNA recognition at many sites in the human genome.

### An algorithm to discover variable-length shape-motifs *de novo* from unaligned genomic regions

We hypothesized that one reason TFs can bind to regions with no evident sequence motifs is their ability to recognize DNA shape independent of the underlying nucleotide sequence. Since different sequences can encode the same values of a DNA structural feature, shape recognition might occur in the absence of sequence motifs. To explore this idea, we first defined the concept of a TF shape-motif, which is a significantly over-represented *pattern* in the profiles of DNA shape features at the TF’s binding sites as compared to non-bound regions (Fig. 1, see Methods). For instance, we would say that a TF has a minor groove width shape-motif if its binding sites are enriched for windows with a particular sequence of minor groove values (e.g., low, high, low) compared to flanking non-peaks. This definition is based directly on the shape feature and is not conditional on the presence of particular nucleotides. It therefore enables us to study shape preferences independent of sequence preferences, which has not been done in previous analyses of DNA shape in the context of TF binding.

**Figure 1.**
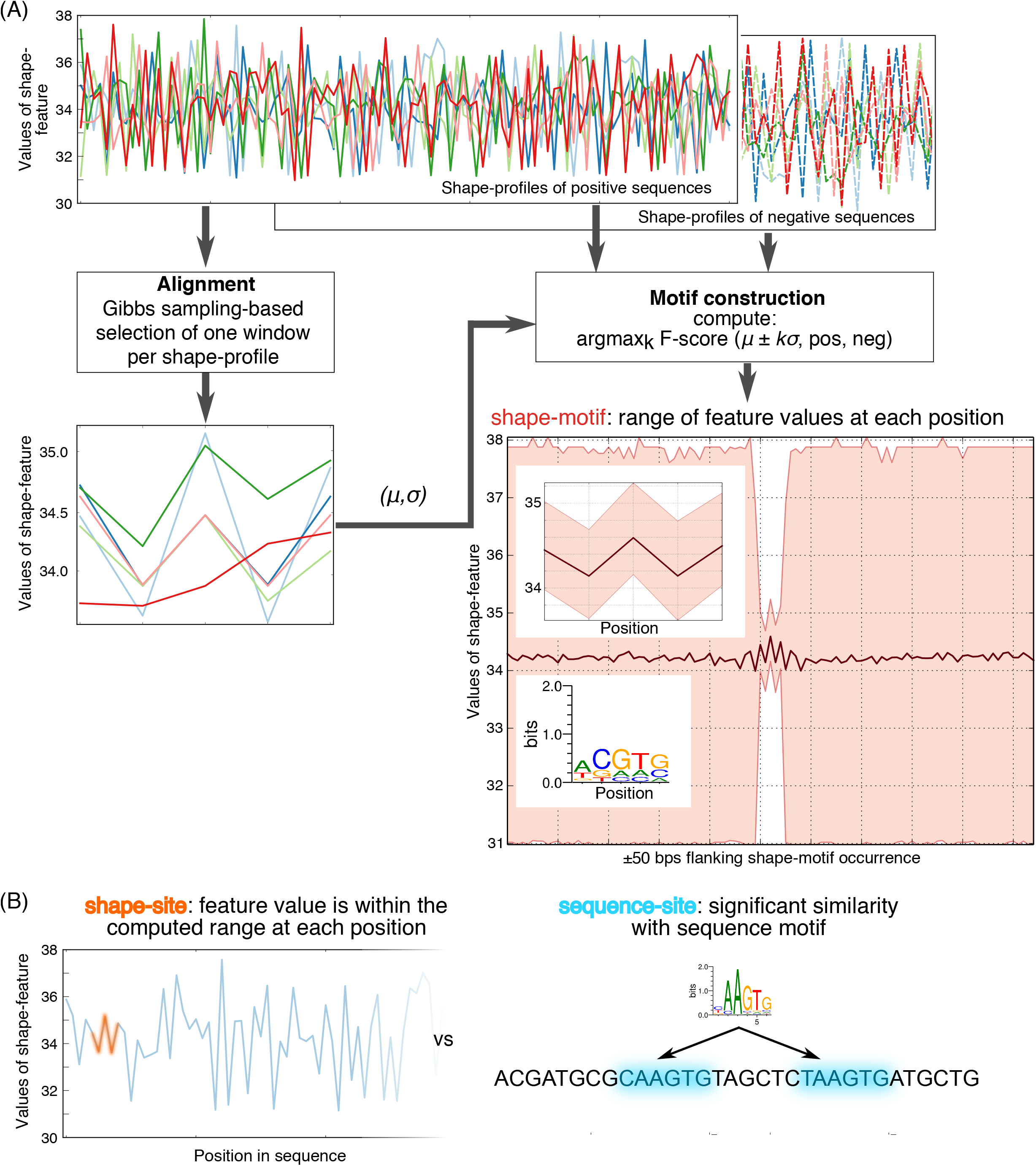
Overview of ShapeMF. (A) Shape-motif discovery involves comparing TF-bound regions (positives; solid lines) to flanking non-bound regions (negatives; dashed lines). For each region (different colors) and each shape feature (MGW, HelT, Roll, ProT), ShapeMF extracts the profile of quantitative feature values across the region. The Gibbs sampler then identifies a set of short windows (5 to <30 bp) from the profiles of positive sequences that have similar patterns of the shape feature. In the second step, this initial set of positive region windows is refined so that the resulting windows share a shape pattern that has the maximum accuracy to discriminate between positives and negatives (Methods). The range of feature values from this refined set of windows defines the shape-motif. For visualization purposes, we also generate sequence logos from the sequences underlying the occurrences of a shape-motif and the range of feature values in the 50-bp regions flanking up- and downstream its occurrences (both shown below the shape-motif). (B) Difference in the approach of identifying a shape-motif occurrence vs. a sequence-motif occurrence. A shape-motif occurs in a sequence if it contains a window whose feature values at every position fit within the ranges defined by the shape-motif. A sequence-motif occurs in a sequence if it contains a window that is significantly similar to the multinomial model defined by the sequence-motif.

Our two-step algorithm to find shape-motifs is called *ShapeMF* (Fig. 1). It extends a typical approach to sequence motif discovery to instead identify profiles of quantitative DNA shape feature values. The resulting shape-motifs retain all the constraints that sequence-motifs satisfy (see Methods). Before searching for shape-motifs, we translate DNA sequences of bound regions and matched unbound regions into a vector of shape features at each nucleotide. The shape features we utilize are helical twist (HelT), minor groove width (MGW), propeller twist (ProT), and roll (Roll), which are estimated from molecular dynamics simulations and available from the GBshape database (Chiu, Yang et al. 2015). These could easily be extended to include additional structural features. Our algorithm operates directly on the quantitative shape values for each site and does not use DNA sequence in any other way. Different nucleotide sequences can encode similar shape, so an enriched shape pattern in bound regions need not correspond to an enriched sequence pattern.

In the first step of the algorithm, we apply Gibbs sampling to compute local alignments of windows from shape-profiles of the bound regions (one per region) without using the unbound regions. The second step uses this alignment to compute a subset of windows whose shape defines a pattern that significantly discriminates bound from unbound regions. Different sets of unbound regions can be used to identify shape-motifs that are discriminative in different contexts. By repeating the search with different window sizes, the algorithm can identify variable-length shape-motifs. The final output includes only the non-redundant motifs.

ShapeMF is implemented in Python and freely available at: https://github.com/h-samee/shape-motif. The software takes as input sets of bound and unbound regions and outputs enriched motif profiles with p-values.

### Most TFs have at least one shape-motif

We applied ShapeMF to discover shape-motifs in the K562 and Gm12878 ChIP-Seq peaks of 110 TFs (201 ChIP-Seq datasets). For each TF in each cell line, we took the strongest 2000 peaks and extracted 100 base-pair (bp) windows centered at the peak-summit (i.e., the location of maximum ChIP intensity) as bound regions and flanking 100-bp sequences sampled 200-bp away as unbound regions (see Methods). We found that >80 of the TFs in each cell-line have a shape-motif (71/84 TFs in K562, 50/63 TFs in Gm12878; Bonferroni-adjusted p-value < 10^-5^; median false positive rate: 0.16-0.19) (Fig. 2A). Few TFs have shape-motifs for all features. However, most TFs have a ProT-motif, while Roll-motifs are much less common (Fig. 2B). We found shape-motifs up to 29 bp long, with an average length of 15 bp (Fig. 2C), which is much longer than a typical sequence motif (6–10 bp) (Stewart, Hannenhalli et al. 2012). In the following analyses, we focus only on the TFs for which we found shape-motifs.

**Figure 2.**
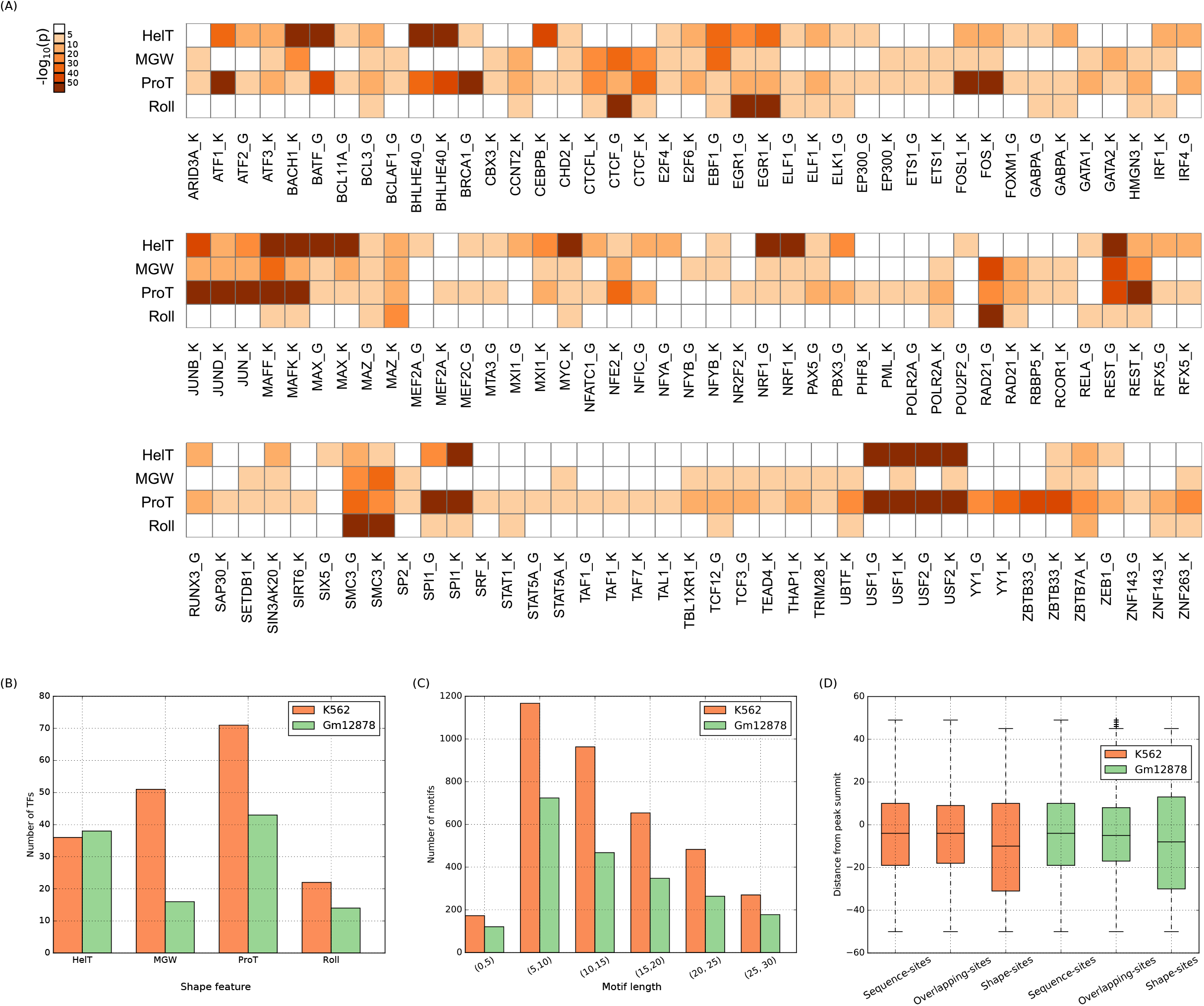
Most TFs have a shape-motif in their ChIP-seq peaks. (A) Heatmap of negative-log10 transformed Bonferroni corrected p-values for the four types of shape-motifs of each TF. White cells indicate no significant motif. ‘K’ or ‘G’ after a TF’s name denotes K562 or Gm12878 cells, respectively. (B) Numbers of TFs with each of the four types of shape-motifs. (C) Length distribution of shape-motifs. (D) Boxplots of relative distances of sequence-, shape-, and overlapping-sites from ChIP-seq peak-summits.

### Shape-motifs are prevalent and distinct from sequence-motifs

We identified all significant sequence matches to each TF’s shape-motifs and sequence-motifs within its genome-wide ChlP-seq peaks in each cell line (see Methods). While we used all five gkmSVM sequence motifs for the TF, we allowed at most one shape-motif per feature, thus using at most four shape-motifs per TF in each cell line. As such, our estimates of shape-motif prevalence are likely conservative. We refer to shape-motif occurrences as *shape-sites,* sequence-motif occurrences as *sequence-sites,* and overlapping occurrences of both types of motifs as *overlapping-sites.*

This analysis identified thousands of shape-sites across the human genome, with a typical TF having 1.56 shape-sites per peak (range: 1.03-5.18 per peak), compared to 2.60 sequence-sites (range: 1.08-4.51), and 3.12 overlapping-sites (range: 1.02-6.61) (Table S2). The higher rate of sequence-sites and overlapping-sites could be driven in part by our being more conservative in calling shape-motifs than sequence-motifs. The amount of TF binding associated with each type of motif is explored below. Both sequence-sites and shape-sites are more prevalent in the top peaks compared to less significant peaks or all called peaks (Fig. S1). Shape-sites (Fig. 2D) generally occur within ±30 bps of the ChlP-Seq peak-summit, as do sequence-sites and overlapping-sites. However, most shape-sites do not overlap a sequence-site (mean: 60.16, range: 16.36-99.16), though overlapping-sites are common for certain TFs, such as USF1, USF2, BHLHE40, CTCFL, EGR1, FOS, and YY1. The GC-content of shape-sites is very similar to that of sequence-sites (66.3 vs. 66.8 in K562, 63.8 vs. 63.4 in Gm12878) and is consistent with a previous analysis of sequence-motif hits in these cell-lines (Wang, Zhuang et al. 2012). Interestingly, overlapping-sites have particularly high GC-content (72 in K562, 69.8 in Gm12878).

Next, we built sequence logos from the shape-sites of each TF and compared these to its sequence-motif logo. This revealed that most shape-motifs are not sequence-specific (Table S3). Their average information content is less than half that of sequence-motifs (6.98 bits versus 15.58 bits for sequence-motifs in TRANSFAC (Matys, Fricke et al. 2003)). Similarly, average shape-motif information content per position is only 0.63 bits, compared to 1.27 bits for TRANSFAC sequence-motifs. The different shape features varied remarkably in their sequence specificity, especially when we consider motifs of different features based on information content per position (ICP): the maximum ICP of an MGW-motif and the minimum ICP of a Roll-motif occur at around the same value of ~0.5 bits, whereas the ICP of HelT-motifs is centered around ~0.5 bits and of ProT-motifs is more uniformly spread within 0.1-1.2 bits (Fig. S2). Across shape features, TFs differed in the information content of their shape-motifs by 3.15-fold. The TFs whose shape-motifs had high sequence information content were largely the same as those with the highest number of overlapping-sites (Fig. S3). Thus, sequence-motifs can encode DNA shape, but shape-motifs frequently occur without a consistent DNA sequence pattern.

### TFs recognize DNA shape of regulatory elements in the absence of sequence motifs

We next examined ChlP-Seq peaks to determine if TF binding at each genomic location is determined by shape-sites, sequence-sites, or a combination. Peaks fall into four categories: (i) *sequence-only,* (ii) *co-occurrence,* (iii) *shape-only,* and (iv) *no motif occurrence* (Fig. 3A). While TFs vary in the proportion of peaks with shape-motifs, shape-recognition is very prevalent. Excluding TFs with no motifs of either type, the majority of peaks for most TFs have a shape-site. For a typical TF, co-occurrence peaks are the most common category, followed by shape-only peaks, with sequence-only peaks being the least common. On average, ~20 of genome-wide peaks in a ChlP-Seq dataset are shape-only. For ~60 and ~50 TFs in the K562 and Gm12878 cell-lines respectively, more than 10 of genome-wide peaks are shape-only (Fig. 3B). The TFs with the most shape-only peaks include EP300, BCL3, MYC, and PAX5. These results demonstrate that most TFs can bind DNA through shape recognition without underlying nucleotide specificity.

**Figure 3.**
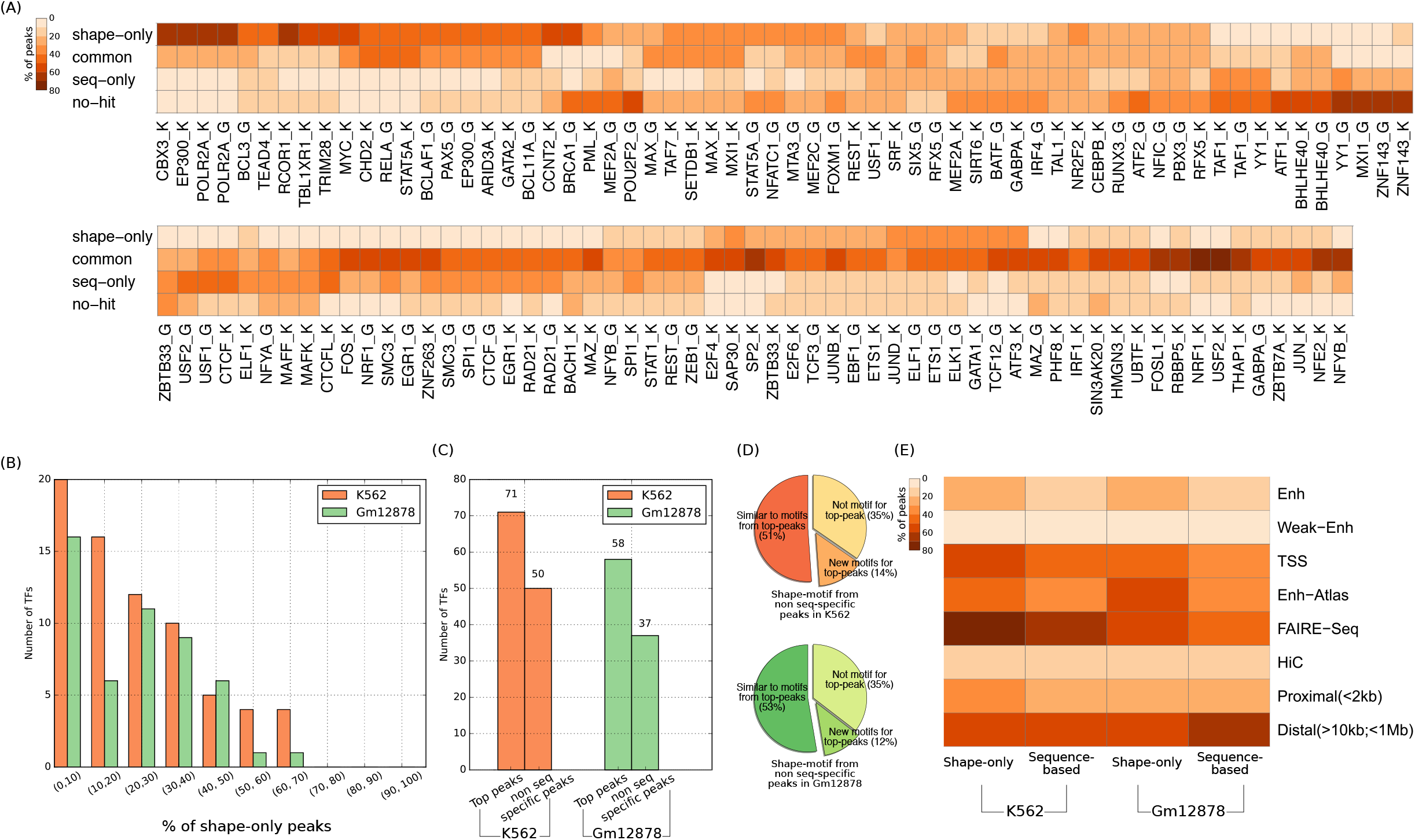
Shape-motifs are common and occur independently of sequence-motifs. (A) The proportion of ChIP-seq peaks that are shape-only, common, and sequence-only for each TF and cell line. (B) Shape-only peaks comprise a considerable fraction of genome-wide peaks for many TFs. (C) For most TFs, shape-motifs can be discovered from the peaks that lack a sequence motif. (D) Most (~65) of these shape-motifs can discriminate between top peaks and their flanking non-peaks, and many (~50) are indeed similar to the shape-motifs discovered from top peaks. (E) Shape-only peaks are generally as abundant as sequence-based (sequence-only and common) peaks in different types of regulatory regions.

To more rigorously test whether shape-specificity is present in the absence of sequence-specificity, we repeated shape-motif discovery on the set of peaks where we could not find any occurrence of a gkmSVM motif. For 58/84 TFs in the K562 cell-line and 37/63 TFs in the Gm12878 cell-line, we found a shape-motif (Fig. 3C). Importantly, 65 of the shape-motifs discovered using peaks without sequence-motifs were also found using the top peaks. Of these, ~80 are similar to the motifs discovered from the top peaks (Fig. 3D). Altogether this analysis supports the conclusion that shape-specificity commonly occurs independent from nucleotide recognition and is a potential explanation to TF binding in regions that lack a sequence-motif.

To explore the functional role of TF shape recognition, we checked how likely a shape-only peak is to occur in putative regulatory regions as compared to a peak that contains a sequence motif *(sequence-based,* i.e., sequence-only or co-occurrence peak). We first considered the active regulatory categories in ENCODE segmentation predictions (ChromHMM and Segway combined) (Ernst and Kellis 2010, Ernst and Kellis 2012, Hoffman, Buske et al. 2012). Shape-only peaks are more likely than sequence-based peaks to occur in enhancers, weak enhancers, and TSS (Fig. 3E). We discovered the same trend upon analyzing enhancers from EnhancerAtlas (Gao, He et al. 2016), ENCODE FAIRE-Seq regions (Consortium 2012), regions in Hi-C contacts (Rao, Huntley et al. 2014), and promoter-proximal and -distal regions (see Methods). These results strongly suggest a role for shape-recognition in functional regulatory elements.

### Shape-specificity is complementary to sequence-specificity

Because co-occurrence peaks are prevalent for most TFs, we sought to understand the relationship between shape-recognition and sequence-recognition within these peaks. We therefore analyzed the spacing of shape-sites and sequence-sites for each TF to find conserved patterns (Fig. 4A-F, Fig. S4; see Methods). Many pairs of shape- and sequence-motifs lack any preference for occurring within, surrounding, or neighboring each other. However, 39 of shape-sites completely contain a sequence-site for the same TF with shape information encoded by and on both sides of the sequence-motif instance (Fig. 4A). Some TFs representative of this trend are EFOS, ATF3, EGR1, ETS1, and ELF1. The converse also occurs, but less frequently: 12 of shape-sites are completely contained within a sequence-site (flanked on both sides by the edges of the sequence-motif) (Fig. 4B,C). Examples include motifs for ATF1, GATA2, MAX, NR2F2, and SPI1. Thus, shape-motifs explain binding to low sequence information content positions within and flanking corresponding sequence-motifs.

**Figure 4.**
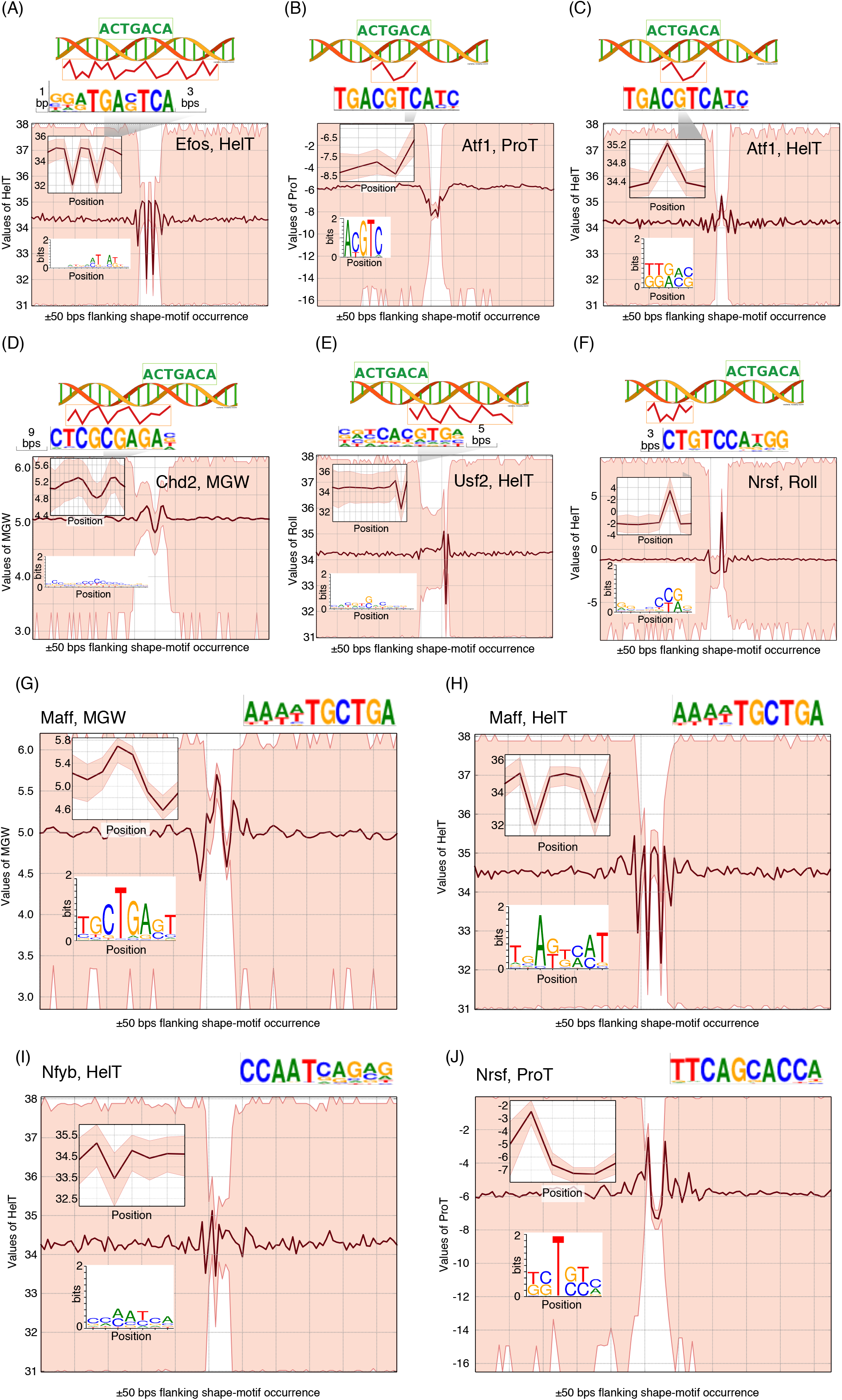
(A-F) Different scenarios of shape- and sequence-motif co-occurrence found enriched in datasets from the K562 cell-line (see Fig. S4 for the same information from the Gm12878 cell-line). Each scenario is shown with a schematic and an example from our analysis. A schematic uses a cartoon DNA double helix (from http://veleta.rosety.com), a sequence `ACTGACA`, and a hypothetical shape pattern to show that a shape-motif can completely contain a sequence-motif (A), a sequence-motif can completely contain a shape-motif (B,C), a shape-motif can overlap with a sequence-motif and flank up- or downstream (D,E), and shape- and sequence-motifs can co-occur with some inter-site spacing (F). Each schematic is accompanied by a real example that includes the sequence-motif (the first of the five sequence-motifs computed by gkmSVM; our analysis uses all five gkmSVM sequence-motifs), the shape-motif, and the inter-site distance that we found enriched in our analysis. We also show the sequence-logo created from sequences underlying the shape-motif, and the range of shape-feature values in the flanking 50-bp regions (up- and downstream) of the shape-site. (G-J) Examples where a shape-motif encodes the binding location of a TF’s dimeri-ation partner.

We also observed many cases of side-by-side sequence-sites and shape-sites. The flanking shape-sites occur both upstream (48 of motif pairs) (Fig. 4D) and downstream (45) (Fig. 4E) of sequence-sites, defined as up to 21 bp away with overlap up to 9 bp. In 2 of motif-pairs, the sequence- and shape-motifs do not overlap (Fig. 4F). These often occur with a conserved inter-motif spacing, which can be up to 16 bp. Examples of TFs with conserved spacing include CTCF, CTCFL, NRSF, MAX, MAFF, and ATF1. Some of these cases correspond to hetero- or homodimer binding motifs. For example, a MAFF MGW-motif encodes a half-site corresponding to the MAFF recognition motif TGCTGA (Yoshida, Ohkumo et al. 2005) (Fig. 4G), and a HelT-motif encodes the TGAGTCA motif of JUN, a well-known co-factor of MAFF (Fig. 4H). Similarly, a HelT-motif for NFYB, commonly located 15 bp upstream of NFYB’s sequence-motif, encodes the same sequence-motif (Fig. 4I). On the other hand, a ProT-motif of NRSF (REST), commonly located 3 bp upstream of the sequence motif of NRSF, does not encode the sequence motif of NRSF, and to our knowledge NRSF does not have any dimerization partner (Fig. 4J). In the following subsection, we analyze this phenomenon of shape-motifs of a TF specifying binding sites for its co-factors in more detail.

Overall, these analyses suggest that shape-specificity is largely complementary to sequence-specificity, and shape- and sequence-motifs collectively define a broader genomic context defining the TF’s putative binding locations. Hence, the shape-specificity of a TF cannot be fully captured by simply taking the shape-profiles underlying sequence-sites. Our results also corroborate and extend the previous findings that implicate flanking nucleotides of sequence-motifs in dictating TF-DNA binding (Dror, Golan et al. 2015) by showing that flanking regions often harbor shape-motifs despite lacking preferred nucleotide patterns.

### Some TFs extensively use shape-specific binding to co-bind with other TFs

Co-binding of TFs is a common phenomenon and is critical for precise spatio-temporal regulation of gene transcription (Gerstein, Kundaje et al. 2012). However, we found that about half (mean 53) of a TF’s peaks that overlap peaks for other TFs *(co-bound* peaks) have no sequence-sites and ~27 are shape-only (Fig. S5, see Methods). We therefore sought to better understand the extent of shape-only binding as a mechanism for TFs to co-bind with other TFs. We found that shape-only binding is more common than sequence-based binding for ~28 of co-bound TF pairs (Fig. 5A, Fig. S6). TFs that mostly use shape-only binding belong to multiple different protein families and include TFs that are known to bind in the context of many other proteins (e.g., MYC, MAX, JUND, STAT5A, GATA2, CCNT2, MEF2A, PAX5, POUF2 (Gerstein, Kundaje et al. 2012)) or to function broadly in genome-wide transcriptional regulation (e.g., EP300).

**Figure 5.**
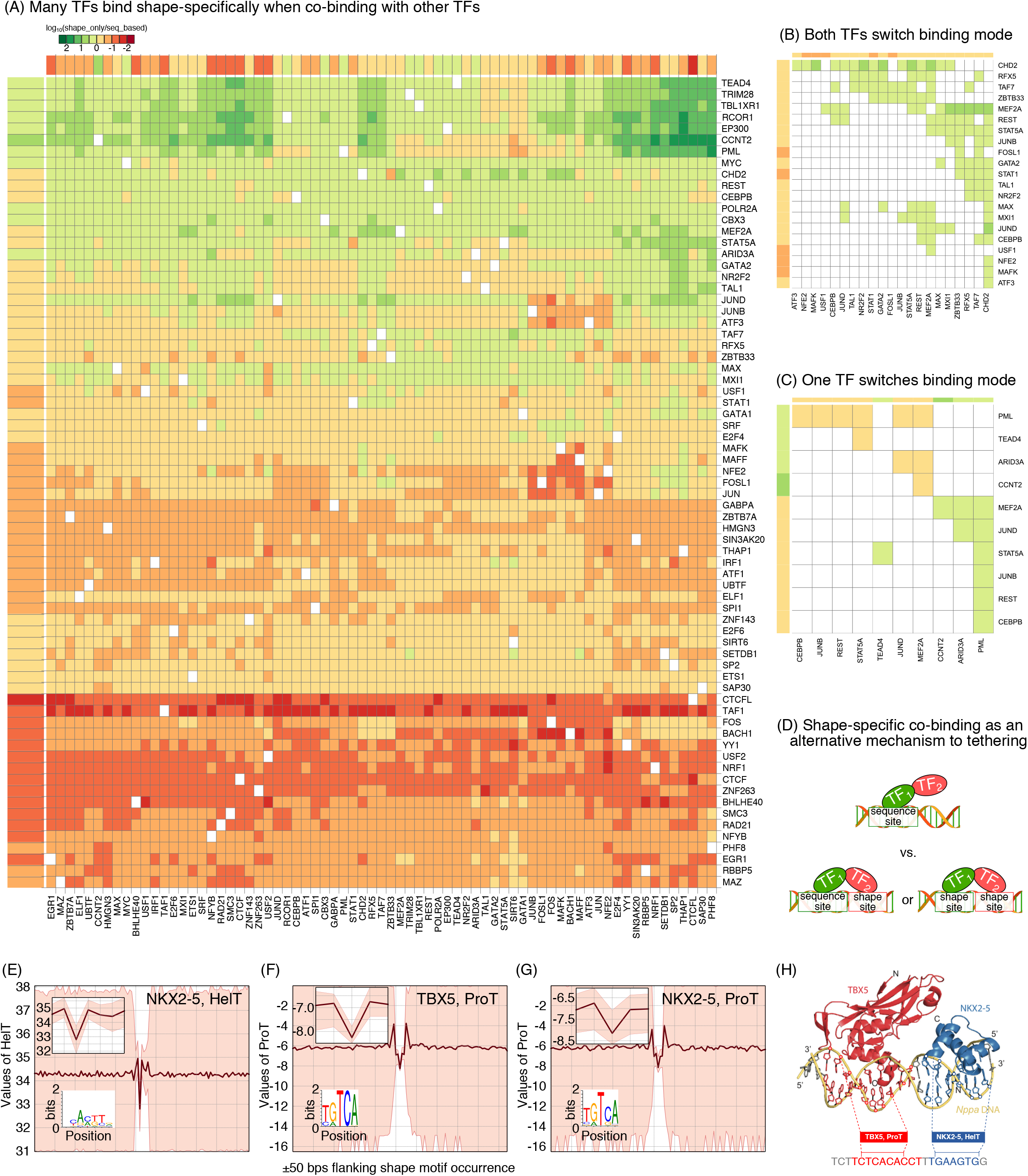
(A) Co-binding TF pairs often utilize shape-specific binding. For each TF pair (f1, f2), the cell at row f1 and column f2 of the heatmap shows whether f1 binds more shape-specifically or sequence-specifically with f2 in the regions where they co-bind in K562 (Gm12878 in Fig. S6). Whether f1’s binding is more shape-specific or sequence-specific (“binding mode”) in the f1-f2 co-binding regions is defined as the number of shape-only peaks being more than that of sequence-based (sequence-only and common) peaks in their co-binding regions. To show whether a TF’s genome-wide binding is more shape-specific or sequence-specific, we use colored bars (following the same scale as in the heatmap) adjacent to the row and the column corresponding to that TF. The colored bars for a TF show whether its genome-wide peaks are more shape-only or sequence-based. The binding mode of a TF in a co-binding region may alter depending on its partner: both (B) or one (C) TF may alter binding mode. (D) Schematic comparing the tethering model with a model of TF co-binding where one or more TFs bind by recognizing DNA shape. (E-G) Shape-motifs of NKX2-5 and TBX5. For each shape-motif, we show the logo created from its underlying sequences and the range of shape-feature values in the flanking 50-bp regions of shape-sites (as in Fig. 4). (H) Occurrences of TBX5 and NKX2-5 shape-motifs in a 22 bp DNA sequence (from mouse *Nppa* promoter) where the two TFs are known to bind. Crystal structure of the ternary complex comprising TBX5, NKX2-5, and DNA is from our previous study.

By examining the prevalence of shape-recognition for each TF in peaks co-bound by every other TF, we learned that TFs use shape- and sequence-recognition differently when binding alone and with each TF partner. First, many TFs enriched for sequence-only peaks genome-wide (e.g., CHD2, REST, CEBPB, MEF2A, STAT5A, GATA2, JUND, MAX, MXI1, MEF2C, MEF2A, POUF2, RUNX3) use mostly shape-sites when co-binding with other TFs. There are also TF pairs where the modes of binding (mostly sequence-based vs. mostly shape-only) of both TFs are different in co-bound peaks compared to the their genome-wide preference (Fig. 5B, Fig. S6). However, the opposite scenario is more common: the same mode of binding is found genome-wide and in co-bound regions (Fig. S7). Also, whether a TF in a given pair will mainly have shape-only binding in their co-binding peaks typically depends on the TF itself. Finally, some TFs that mainly utilize sequence-recognition switch to shape-recognition being more dominant in regions co-bound with a TF that is mainly shape-based and vice versa (Fig. 5C, Fig. S6). We conclude that while the shape preferences of a TF are fairly consistent across the genome, there are many TF pairs where co-binding is characterized by unique shape-motifs.

Co-bound TFs may interact physically and form a complex. In such cases, it is likely that motifs of the partnering TFs occur with some bias in their inter-site spacing. Our observation that co-bound TFs commonly use shape-specific binding led us to hypothesize that some TFs of a DNA-binding TF-complex may bind DNA shape-specifically. This scenario is in contrast with the general notion of “tethering” whereby it is assumed that one or more TFs of a complex recognize the DNA by sequence-specificity and the other TFs do not recognize DNA (Fig. 5D). To assess whether and to what extent TFs that lack sequence-sites in co-bound regions use shape-sites versus tethering, we first evaluated motif spacing and found that 2633 out of 3710 (71) TF pairs have sites that occur with a bias for short (~3 bps) inter-site spacing (Fig. S8). This high proportion raises the possibility that these TFs might form TF-complexes. Interestingly, for 1245 of these pairs, motifs of one or both TFs are shape-motifs. These shape-sequence or shape-shape motif pairs confirm that about a third of adjacent co-binding occurs with at least one TF using shape recognition without nucleotide recognition. We also found that 57 of these pairs have previously been reported to have physical interaction (Stark, Breitkreutz et al. 2006, Ravasi, Suzuki et al. 2010) or predicted for tethered binding (Wang, Zhuang et al. 2012). Some shape-shape motif pairs (e.g., JUN-TAF7, CTCF-YY1, ETS1-TAL1) were not reported in the above studies but were validated elsewhere to have physical interaction or to co-bind (Munz, Psichari et al. 2003, Palii, Perez-Iratxeta et al. 2011, Schwalie, Ward et al. 2013). Therefore, we find an interesting line of evidence that some TFs in a DNA-binding complex may actually bind DNA in a shape-specific manner, and it is unlikely that tethered binding is the only explanation for TF complex members that lack sequence-motifs.

In a ChIP-exo based study of TBX5 and NKX2-5 occupancy in cardiac differentiation, we determined that the binding of these two TFs can be interdependent (Luna-Zurita, Stirnimann et al. 2016). Sequence motifs of TBX5 and NKX2-5 co-occur in only 17 of the regions where their ChIP-exo peaks overlap. Motivated by the above analysis, we hypothesized that TBX5 and NKX2-5 may bind shape-specifically in their co-binding regions. We applied ShapeMF on TBX5 and NKX2-5 ChIP-exo peaks and found that both TFs have shape-motifs for all four features. Importantly, we found strong relationships between the sequence and shape motifs of these TFs, which gave us a strong premise for the above hypothesis. In particular, we found that the sequences underlying the HelT motif of NKX2-5 contain the TF’s sequence motif, CACTT (Fig. 5E). Likewise, the sequences underlying the ProT motif of TBX5 contain a partial sequence motif of TBX5, TGTCA (Fig. 5F). Interestingly, we also found that the sequences underlying the ProT motif of NKX2-5 contain TGTCA sequences (Fig. 5G)-implicating that in many NKX2-5 peaks, TBX5 may co-bind by recognizing the ProT pattern, without full sequence recognition.

In support of our hypothesis that TBX5 and NKX2-5 may bind shape-specifically in their co-binding regions, we found that 79 of co-binding regions have a shape-site for one TF and a sequence motif for the other TF, and 73 contain shape-motifs of both TFs. Furthermore, co-occurrences of the shape- and sequence-motif pairs of TBX5 and NKX2-5 are enriched for 0-4 bps inter-site spacing, which is in the same range as the preferential distances between TBX5 and NKX2-5 sequence motifs that we identified previously, supported by crystal structure (Fig. 5H) (Luna-Zurita, Stirnimann et al. 2016). Overall, this analysis shows that the TBX5-NKX2-5 co-occupancy occurs to a large extent by recognizing DNA shape. The use of shape-recognition in TBX5-NKX2-5 co-bound regions exemplifies the use of shape-motifs as an alternative to tethering, and is highly relevant to coregulation of a cardiac transcriptional program, as well as a potential mechanism for their haploinsufficiency in congenital heart disease.

### Shape-motifs explain genomic occupancy of TF-complexes where sequence-based models are inadequate

The above cases of shape-specific binding in regions of TF-TF co-occupancy motivated us to examine whether DNA shape may explain the genomic occupancy of TF complexes for which sequence-based models have not been able to explain complex patterns of co-binding. We focused on two well-known TF-complexes, namely the MAX homodimer and the MYC-MAX heterodimer, several aspects of whose *in vivo* occupancies have remained unresolved in sequence-based analyses (Guo, Li et al. 2014, He, Johnston et al. 2015).

The bHLHZip domain protein MAX recognizes the canonical E-box motif CACGTG. Current models suggest that MAX can bind DNA either as a homodimer or upon forming heterodimers with other bHLHZip proteins, such as MYC and MAD. Max footprints in ChIP-nexus data (He, Johnston et al. 2015) were found to span ~8 bases flanking either side of the E-box motif. Although the footprints were enriched for the E-box motif, the footprints covered the E-box motif only partially and there was no specificity in the flanking sequences. Our analysis of MAX ChIP-Seq peaks identified HelT-, MGW-, and ProT-motifs for Max, and suggest an interesting model involving DNA shape and sequence determining the specificity of MAX homodimers (Fig. 6A). In particular, sequences underlying the HelT-motif directly match the E-box sequence motif (Fig. 6B, left panel). On the other hand, the MGW- and ProT-motifs show very low sequence-specificity and in the co-occurrence peaks (i.e., where these shape-motifs co-occur with MAX’s sequence motif), they occur 6-10 bases up- and 4-5 bases downstream of the E-box motif, respectively (Fig. 6B, middle and right panels). This result provides a shape-based explanation of MAX binding where specificities for HelT, MGW, and ProT explain Max binding both at the E-box motif and the inclusion of flanking sequences in Max-bound footprints.

**Figure 6.**
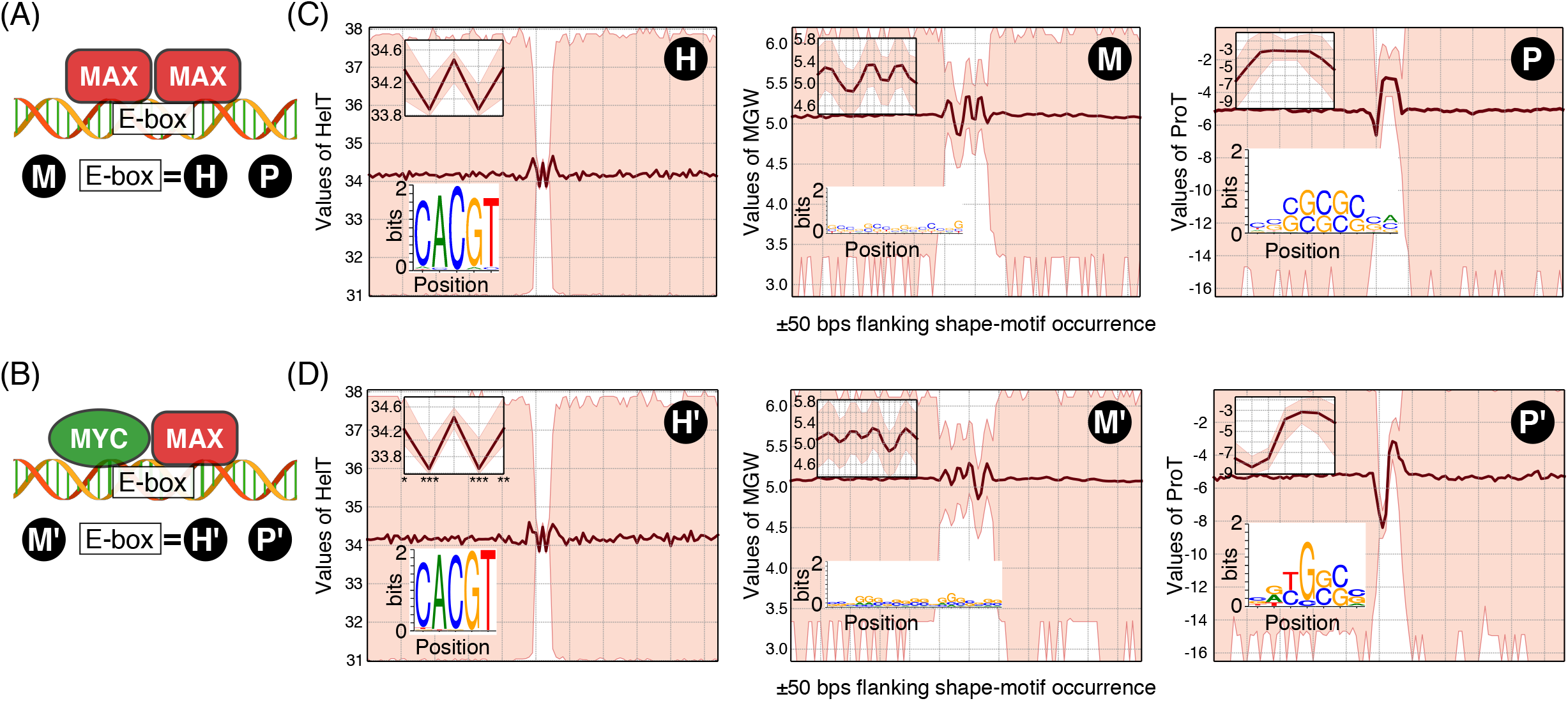
Shape-motifs suggest models for genomic occupancy of (A) MAX homodimer and (B) MYC-MAX heterodimer. The HelT-, MGW-, and ProT-motifs enriched under (C) MAX ChIP-Seq peaks vs. negative controls and (D) MYC vs. MYC-unbound MAX-peaks. For each shape motif, we show the logo created from its underlying sequences and flanking 50-bp regions (as in Fig 4). HelT values in the MYC-MAX motif (left panel in D) differ significantly from the MAX HelT motif (left panel in C) in positions 1 (Kolgomorov-Smirnov test p-value < 0.05), 2 (p-value<0.005), 4 (p-value < 0.005), and 5 (p-value < 0.01).

MYC is another bHLHZip domain protein known to bind very weakly at E-box motifs as a monomer, but binds the same sequences with high affinity upon dimerization with MAX (Guo, Li et al. 2014). Our analysis of MYC K562 ChIP-Seq data supports this model, since ~75 MYC peaks overlap MAX peaks, and the intensity of MYC ChIP signal has a strong dependence on the extent of overlap (Fig. S9, see Methods). However, MAX binds in almost twice as many locations as MYC, and it is not clear how the specificity of the MYC-MAX dimer is different from that of MAX in the other MAX-bound locations. There is a hypothesis that bases flanking E-box motifs play a role in MYC-MAX co-bound regions (Nair and Burley 2003).

We hypothesized that MYC-MAX binding could be distinguishable from the binding of MAX homodimers in terms of shape-specificity. We therefore performed differential shape-motif discovery from MYC-MAX peaks utilizing MYC-unbound MAX regions as the negative control. This analysis indeed suggests a model whereby MYC-MAX binding is distinct from the binding of MAX homodimers in terms of DNA shape (Fig. 6C). We identified HelT-, MGW-, and ProT-motifs for the MYC-MAX dimer and found that the HelT-motif for MYC-MAX encodes the E-box sequence motif (Fig. 6D, left panel), similar to the case for the MAX homodimer. Interestingly, HelT values at certain positions of the two motifs are significantly different (Fig. 6D, left panel, positions shown with asterisks). Moreover, the MGW- and the ProT-motifs for MYC-MAX are different from those of MAX in terms of length, pattern, the sequences underlying those motifs, and their spacing with the E-box motif (Fig. 6D, middle and right panels). In the co-occurrence peaks, the MGW-motif is often located 12-14 bp upstream of the E-box motif and the ProT-motif occurs 1-3 bp downstream. It is known that crystallized structures of the MYC-MAX dimer and the MAX homodimer are different despite their apparent resemblances (Nair and Burley 2003). Combining this with our shape-motif analysis, we speculate that the structural differences between the MYC-MAX dimer and the MAX homodimer cause subtle differences in their DNA-binding specificities that are largely accommodated by changes in MGW and ProT profiles. Overall, the above examples suggest that TF complexes, like individual TFs, utilize shape recognition as a mechanism to discern their binding locations.

### Non-targeted TF motifs (aka “zingers”) can be shape-specific

Hunt and Wasserman recently reported *zinger* motifs: sequence-motifs of a small group of TFs enriched across the binding locations of multiple other TFs (Worsley Hunt and Wasserman 2014). Along the same lines, we next asked if there are *shape-zingers,* i.e., shape-motifs enriched across ChIP-Seq peaks for many TFs. We tested for enrichment of shape-sites for each TF within the top 2000 peaks of every other TF. Results were consistent when we repeated the analyses using all peaks without a sequence-site for the other TF. We found that ~25 ENCODE TFs are shape-zingers, including previously reported sequence-zingers, such as JUN, FOSL1, and the ETS-family TFs EBF1 and ELK1. However, most shape-zingers are not sequence-zingers (Fig. 7, Fig. S10). Importantly, some of these novel shape-based zingers (e.g., GATA, ARID3A, P300, PAX5) are known to act as global regulators or regulators of large gene networks (Goodman and Smolik 2000, Medvedovic, Ebert et al. 2011, Rhee, Lee et al. 2014, Lentjes, Niessen et al. 2016). This finding suggests that shape-specificity enables these regulators to recognize a larger set of locations (and thus regulate more genes) than would be possible based on sequence-specificity alone. Consistent with the “loading station” model suggested by Hunt and Wasserman, all our shape-zingers (except P300 in the Gm12878 cell-line) show enrichment within peaks of CTCF, RAD21, and SMC3.

**Figure 7.**
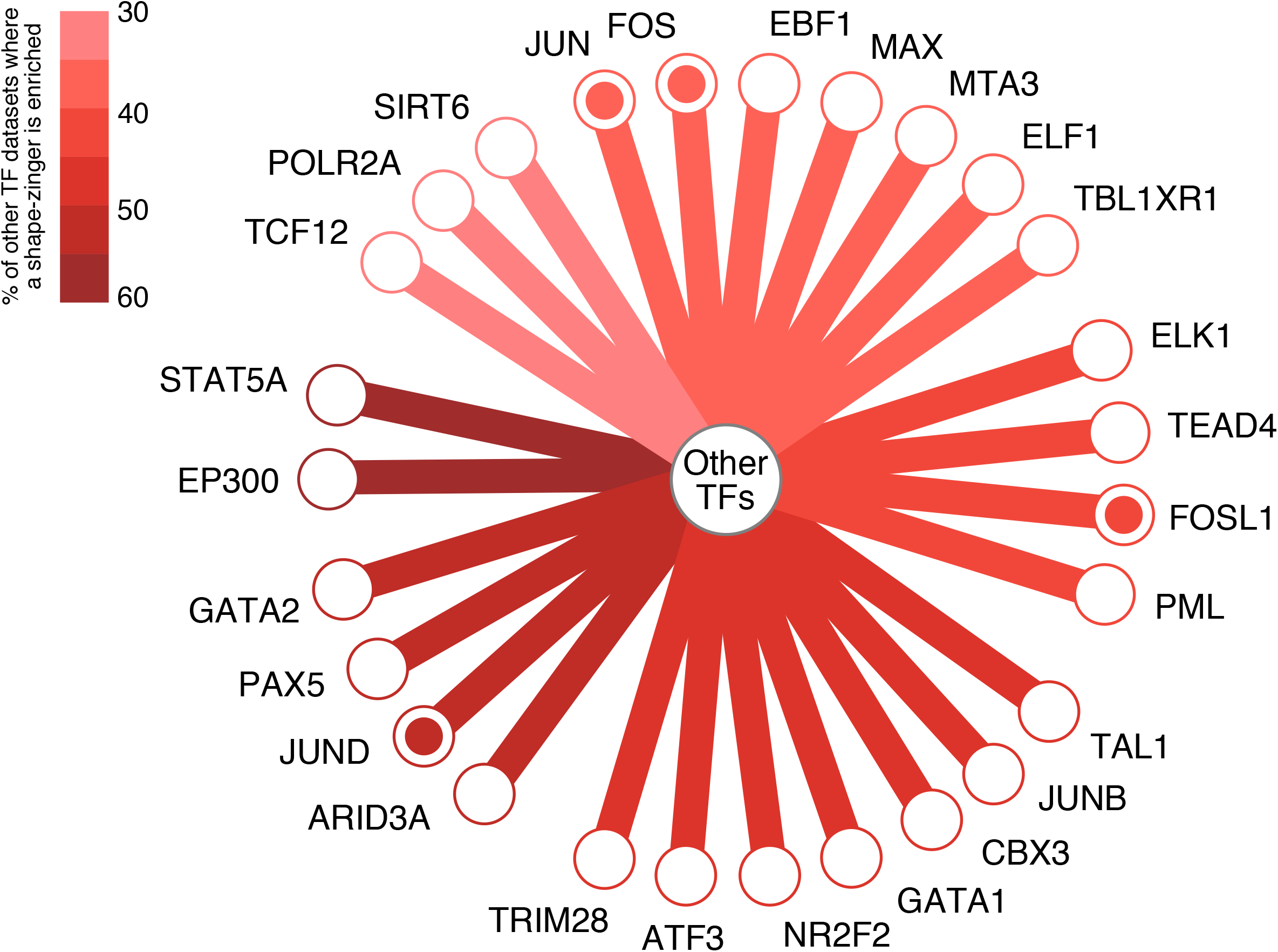
Shape-zinger TFs (nodes) and the fraction (color-coded) of other ChIP-Seq datasets where their shape-motifs are enriched in. Shape-zingers are the TFs with shape-motifs enriched in the largest numbers (at least 30, see Methods) of other TF’s ChIP-seq datasets. Enrichment was calculated in the top 2000 peaks vs. non-peaks (Methods). TFs corresponding to nodes with a filled circle were previously reported as sequence-zingers. For brevity, edges connecting shape-zinger pairs are omitted.

### TFs within the same class recognize distinct shape-motifs

TFs within the same class of DNA-binding domain often recognize statistically indistinguishable sequence motifs, although such TFs still bind distinct locations in the genome. It has been shown that sequence-based models of TF-occupancy achieve improved performance for TFs within the same class if the models use shape-features underlying sequence-motif hits (Mathelier, Xin et al. 2016, Yang, Orenstein et al. 2017). However, it is not clear whether TFs within a class show preferences for distinct shape-patterns and/or for distinct combinations of shape-features. To test this possibility, we applied ShapeMF on ENCODE ChIP-Seq datasets of the bHLH and bZIP class TFs (see Methods). For each TF within a class, we considered only those ChIP-peaks that do not overlap with peaks of any other TF within that class so that we could identify the shape-specificity intrinsic to each TF. Our analysis not only found that most TFs within these two classes recognize distinct motifs of different shape features, but also that some shape-motifs do not encode the common sequence-motif of that class (Fig. 8, Fig. S11). Overall, we conclude that preference for distinct shape-patterns, and sometimes for distinct combinations of shape-features, is a mechanistic explanation of how TFs within the same class of DNA-binding domains bind at distinct locations genome-wide.

**Figure 8.**
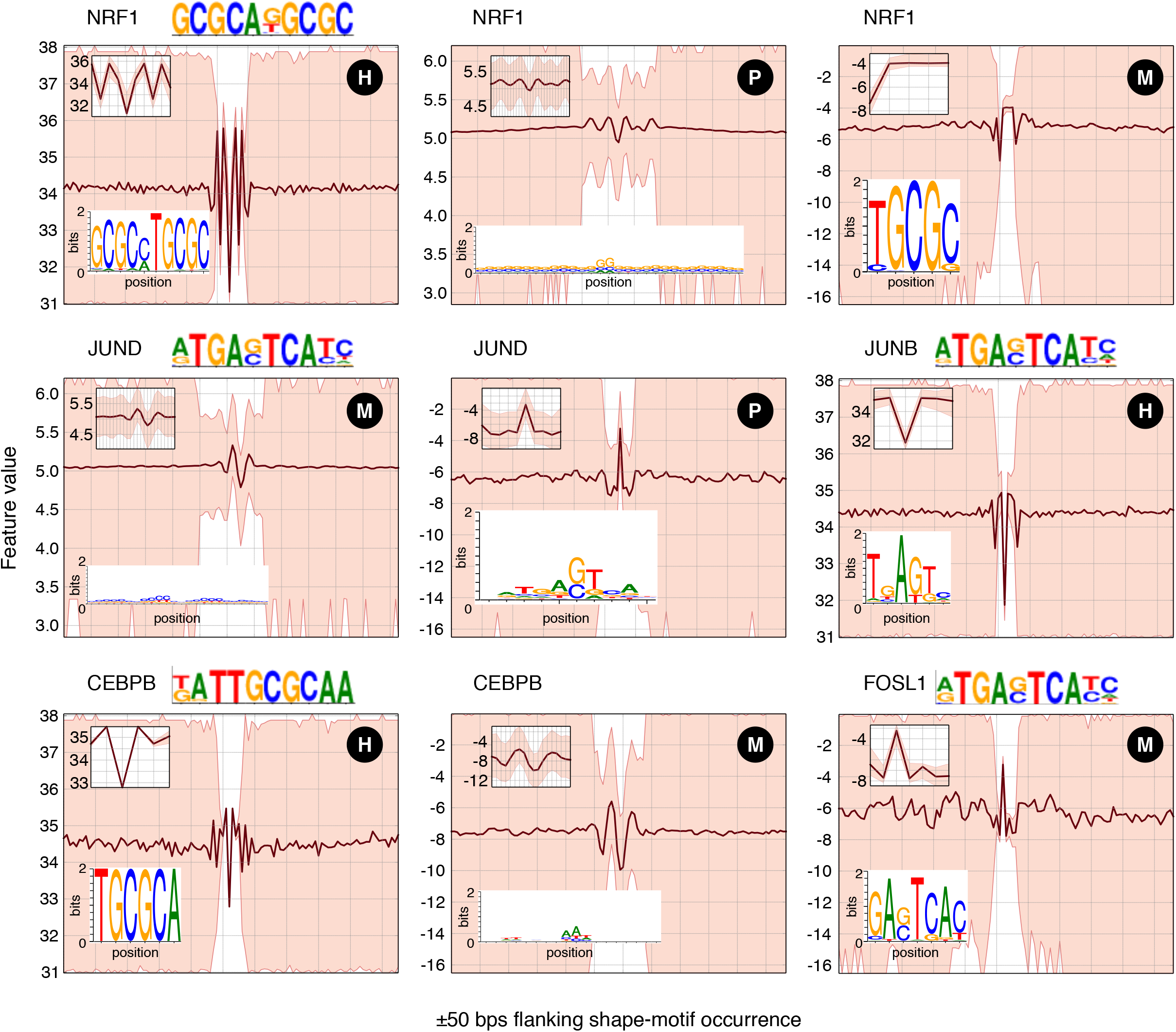
bZIP TFs have similar sequence-motifs but unique shape-motifs. Shape-motifs for five bZIP proteins NRF1, JUND, JUNB, CEBPB, and FOSL1 are shown (H, M, and P denote motifs for HelT, MGW, and ProT, respectively). A feature is not mentioned for a TF if the TF does not have a shape-motif for that feature. For each TF, the first of its five gkmSVM motifs is shown to aid comparison with the logos created from sequences underlying the TF’s shape-motifs.

### TFs bind shape motifs *in vitro*

To complement and validate our analyses of shape-motifs in ChIP-Seq peaks, we analyzed their occurrence in oligonucleotides that have been assayed for *in vitro* binding. Nineteen ENCODE TFs were investigated via HT-SELEX (Jolma, Yan et al. 2013, Yang, Orenstein et al. 2017) and have shape-motifs that are 15-bp or less, which is short enough to be present in these libraries. We found that shape-motifs are enriched in the final round (i.e., preferentially bound) oligonucleotides for all 19 TFs. Furthermore, shape-motif prevalence is in good agreement between HT-SELEX and ChIP-seq data (correlation 0.54; Fig. S12). We additionally found enrichment of shape-motifs for MAX and MYC amongst bound versus unbound oligonucleotides in genomic-context protein binding microarray (gcPBM) data (Gordan, Shen et al. 2013, Afek, Schipper et al. 2014). Shape-motifs of MAX and MYC frequently co-occur with their sequence-motifs, as expected since these gcPBMs were designed to contain sequence-motifs (Zhou, Shen et al. 2015). Thus, despite several design features that bias currently available *in vitro* data against shape-specific binding (see Discussion), we observe a strong and consistent signal that TFs bind shape-motifs when they do occur in assayed oligonucleotides.

## Discussion

Analyzing *in vivo* binding data of hundreds of human TFs with a novel algorithm that treats DNA as a structure rather than a string of nucleotides, we showed that TFs frequently bind DNA by recognizing specific patterns of DNA shape features. These shape-motifs can occur independently from the TF’s nucleotide sequence-motifs, and ChIP-Seq peaks that contain a shape-motif but no sequence-motifs are as abundant as peaks with only sequence-motifs or with both. Shape motifs also shed light on TFs that bind low information-content sites and weak matches to sequence-motifs. Thus, in addition to confirming the importance of DNA shape in the context of sequence-motifs as shown in several recent studies (Abe, Dror et al. 2015, Zhou, Shen et al. 2015, Mathelier, Xin et al. 2016), our results establish shape-recognition as an independent and sometimes exclusive mechanism for TF-DNA binding within regulatory regions.

Our analyses also reveal important functional and mechanistic consequences of shape-based TF-DNA binding. Binding of TFs within the same family to distinct instances of a shared sequence-motif can be explained by TF-specific shape-motifs that overlap or flank the sequence-motif. Similarly, TFs (e.g., MAX) that bind different instances of a sequence motif as monomers, homodimers, or heterodimers appear to recognize different DNA shapes in each of these contexts. These examples suggest that DNA shape may be a general mechanism to increase the information content of binding sites beyond that encoded by sequence-motifs, which is insufficient for eukaryotic TFs to uniquely recognize specific sites in genomic DNA (Wunderlich and Mirny 2009). We also find that co-binding TF pairs frequently utilize shape-based binding, providing a mechanism beyond tethering to explain co-binding in regions that lack one or both sequence-motifs. Finally, TF’s in crystal structures generally contact nucleotide bases at sequence-specific binding sites and the DNA backbone at non-sequence-specific sites (Aishima and Wolberger 2003, von Hippel 2004, Romanuka, Folkers et al. 2009). Our results suggest that shape-specific sites contain preferential “pockets” – defined by shape-motifs – where a TF can stabilize and interact with the DNA backbone. It is also plausible that such stabilization is facilitated by enhanced electrostatic potential at the location of shape-motif occurrences (Rohs, Jin et al. 2010). Importantly, these new insights would have been missed if we had only searched for shape patterns within the context of sequence-motifs.

An important methodological contribution of our manuscript is ShapeMF, a *de novo* shape-motif discovery algorithm. ShapeMF enabled us to pursue the hypothesis that some TFs have intrinsic preferences for shape-motifs and such preferences can be discovered without taking sequence information into account. It is challenging to design such an algorithm since it requires discovering variable-length shape-patterns *de novo* from unaligned sequences with the criterion that the discovered shape-motifs are comparable to sequence motifs in terms of discriminating bound from unbound regions. Our solution was to implement Gibbs sampling with a notion of similarity between the shapes of two DNA sequences that is appropriate for quantitative features, rather than the discrete four-letter nucleotide alphabet. We considered several alternative solutions that have been used on related problems. For example, we might have discretized shape features and then directly applied *de novo* sequence motif algorithms, as in (Greenbaum, Parker et al. 2007). However, it was not clear how to appropriately bin and/or smooth shape features, or how to characterize the background distribution for these features. Time series “shapelet” discovery algorithms are relevant to our problem (Ye and Keogh 2009, Grabocka, Schilling et al. 2014, Hou, Kwok et al. 2016), but their efficacy has been shown for datasets where discriminative shapelets appear very frequently (so that sampling a very small subset of the data would suffice to yield shape-motifs) or where the constituent time series are aligned. These scenarios do not hold for TF occupancy data, so shapelet algorithms would likely require a computationally intensive brute-force scheme. In contrast, ShapeMF does not (a) discretize data, (b) use any empirical background distribution of shape features, nor (c) assume that the given peaks/windows are aligned.

The focus of our study was to systematically test if DNA shape provides signals of “intrinsic” TF-DNA binding specificity (Stormo and Zhao 2010) and if these are independent of nucleotide sequence. We therefore did not adopt the approach of developing an optimal, holistic classifier of bound versus unbound regions. State of the art approaches to this classification problem perform well and can score a given sequence for its affinity. But a method such as ShapeMF is needed to discover the specificity signal that characterizes a TF (Setty and Leslie 2015). Adding shape-motifs to discriminative classifiers of TF bound regions does have potential to improve accuracy, since we found that many peaks with shape-motifs lack sequence-motifs and unbound regions with sequence-motifs can lack shape-motifs.

The intrinsic binding specificity of shape-motifs should be comprehensively studied *in vitro.* We conducted an initial evaluation using HT-SELEX and gcPBM data and found consistent evidence for *in vitro* binding. However, current *in vitro* data have some limitations for evaluating shape-motifs and disentangling the contributions of shape versus nucleotide recognition. Critically, existing oligonucleotide libraries do not contain sufficient coverage of shape-motifs. For example, 86 of the HT-SELEX oligonucleotides we analyzed are 20 bps or shorter (Jolma, Yan et al. 2013, Yang, Orenstein et al. 2017), and universal PBMs (uPBMs) that are designed to compactly cover all k-mers typically have values of k=8 or 10 bp (Berger and Bulyk 2009). Relatedly, HT-SELEX and uPBM oligonucleotides do not fully capture genomic context that may be important to shape-specific binding. These limitations could be overcome with improvements in the throughput of these technologies to accommodate more, longer sequences or by designing gcPBM libraries to contain oligonucleotides with better representation of shape-motif containing genomic regions, including more regions that contain shape-motifs but no sequence-motifs. It will be important to quantify binding affinities for many TFs and shape-motifs with these and other emerging technologies (e.g., microfluidics).

Several other future directions are suggested from our results. One goal is to quantify the amount of information that each TF utilizes from the shape domain and how specific this utilization of DNA shape is to different contexts, including chromatin domains, co-binding, and subsets of target genes. Another important direction will be to re-analyze *in vitro* DNA-binding data to assess whether high-affinity sequences are enriched for shape-based binding. Additionally, the mechanisms of shape-based binding suggested by our results should be evaluated in light of all available TF-DNA structures. It will also be interesting to determine the extent to which shape-based binding is an alternative explanation to tethering. In terms of methodology, we could likely improve the performance of ShapeMF by adding more shape features and/or analyzing these jointly rather than individually. Our current analysis suggests that multivariate analysis of shape-features would require careful algorithm design since a TF may not have a motif for every feature and the motifs may differ in size and location. Finally, the notion that shape-based TF binding affinity could be conserved without sequence conservation opens the door to a whole new view of regulatory evolution and the opportunity to develop shape aware measures of DNA change and its functional impact on human disease that has at its basis abnormal TF function.

## Data and Methods

### The ShapeMF tool for *de novo* shape motif discovery from shape-data

#### Definitions and notation

Let 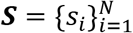 denote a set of peaks of a TF *T*. Let 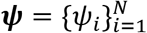 denote the corresponding shape-data for a feature *F* (roll, helical twist, propeller twist, or minor groove width in the current GBshape-based implementation). Thus each *ψ_i_* is a sequence of real numbers *ψ_i,j_* denoting the value of feature *F* at position *j* in peak *s_i_*. A shape *pattern* of length *l* is a *l*-iength sequence 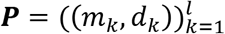 of 2-tuples of real numbers. We say that ***P** occurs* in peak *s_i_* if there is a *l*-length sequence window 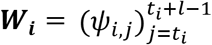 starting at position *t_i_* in the shape-data *ψ_i_* such that *m_j−t_i__*+1 – *d_j−t_i__*+1 ≤ *ψ_i,j−t_i__*+1 ≤ *m_j−t_i__*+1 + *d_j−t_i__*+1 for *t_i_* ≤ *j* ≤ *t_i_* + *l* – 1.

#### Algorithm

ShapeMF uses a two-step approach to first compute a shape pattern **P** from the shape-data of positive peaks in a training dataset of matched positive and negative control peaks and then modify the pattern to one that maximizes F-score between the positive and control peaks in the training data. Finally, the modified pattern is called a motif if its Bonferroni-corrected hypergeometric p-value, computed on a separate validation dataset of matched positive and control peaks, is significant.

**Step 1.** From the shape-data ***ψ*** of positive peaks in the training data, we first compute a set of windows ***W*** (one window ***W_i_*** in each *ψ_i_*) such that the sum of pairwise Euclidean distances between the windows, *i.e.*, *D* = ∑_*x≠y*_ Euclidean(***W_x_,W_y_***), is minimized. We use Gibbs sampling to compute such a set of windows. In particular, we start by selecting ***W_i_***’s randomly. To select a new window from *ψ_i_* to replace ***W_i_***, let 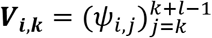 denote the *l*-length window in *ψ_i_* that starts at position *k*, and let
*D_i,k_* = ∑_*x≠y≠i*_ Euclidean(***W_x_W_y_***) + ∑_*j≠i*_ Euclidean(*W_j_,V_i,k_*) denote the new value of *D* if ***V_i,k_*** replaces ***W_j_*** in the current set of windows. We then sample a window ***V_i,k_*** from *ψ_i_* with probability 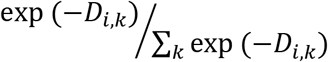 and update ***W_i_*** with the sampled ***V_i,k_***. We iterate through the *ψ_i_*’s in the order they appear in ***ψ***, and for each *ψ_i_*, we update the window ***W_j_*** following the above steps. We continue to repeat this iterative updating until convergence in the value of *D*. In our implementation, we decide that the value of *D* has converged if it does not change in two successive iterations, or if the improvement in the value of *D* is negligible for ten consecutive iterations. We assume that the improvement is negligible when the current and the previous values of *D* satisfies: *D_prev_*(1 – *∈*) < *D_current_* < *D_prev_*, where *∈* = 10^−5^.

**Step 2.** We next compute a pattern 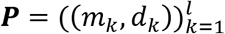 from the set ***W*** of windows computed above. Let the window ***W_i_*** start at the position *t_i_* in *ψ_i_*. We then take *m_k_* = mean (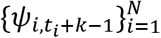) and *d_k_* = *α* × standard_deviation (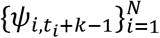), where *α* is a constant whose optimum value is computed as follows. We try each value of *α* in the range [0.1, 2], in increments of 0.1, and compute the corresponding pattern ***P_α_*** from ***W***. We quantify the goodness of each ***P_α_*** as a discriminator between the positive and control peaks of the training data by its *F*_1/3_-score. Note that the *F*_1/3_-score here is a more conservative objective than used typically in classification settings, yet we wanted to weigh precision much higher than recall so that the number of false positives remains low. We then choose ***P*** to be the pattern argmax_α_ (*F*_1/3_(**P_α_**)).

#### Motif identification

Using an independent validation set of positive and control peaks, patterns are tested for enrichment in positive peaks, as is done in other discriminative motif finding tools (Bailey 2011). Let 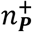 and 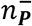 denote the number of validation positive and control peaks, respectively, where pattern ***P*** has an occurrence. We say that ***P*** is a *motif* of feature *F* (or a *‘F*-motif’) for TF *T* if a hypergeometric test parameterized by 2*N, N*, 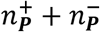 and 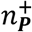 yields a significant *p*-value after Bonferroni correction. We use a Bonferroni corrected *p*-value threshold of 10^−5^. Shape-motifs ***P*** that meet this criterion are retained for further analysis, and others are discarded.

#### Finding variable-length motifs

The above steps 1 and 2 compute a motif for a given length *l*. In our analysis we have considered all values of *l* between 5 and 30. For computational efficiency, we first compute motifs for values of *l* that are multiples of 5. For all other values of *l*, we take the starting positions *t_i_*’s of the motifs computed for length 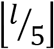 as our initial guess for starting positions and search for motifs within the positions *t_i_* – *l* and *t_i_* + *l*.

#### Calling redundant motifs

In the last step, we eliminate redundant motifs. We first partition all motifs according to their lengths: two motifs of lengths *l*_1_ and *l*_2_ are put in the same partition if 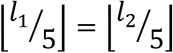. For two motifs ***P*_1_** and ***P*_1_**, if ***P*_1_** has lower false positive rate than ***P*_2_** on validation data and the occurrences of ***P*_1_** “cover” at least 75 of the occurrences of ***P*_2_**, then we assume that ***P*_2_** is redundant and discard ***P*_2_**. An occurrence of ***P*_1_** *covers* an occurrence of ***P*_2_** if the occurrence of ***P*_1_** overlaps with at least 75 length of the occurrence of ***P*_2_**. This strategy to remove redundant motifs is akin to the one utilized in (Beer and Tavazoie 2004).

### Data

We downloaded uniformly processed ChIP-Seq datasets (narrowPeak format) from the *ENCODE Downloads* section at UCSC (<u>http://genome.ucsc.edu/ENCODE/downloads.html</u>) and genome-wide shape-data (bigwig format) from the FTP interface of the GBshape database (<u>http://rohsdb.cmb.usc.edu/GBshape/</u>). We used only the ChIP-Seq datasets with `quality=good` and `treatment=None`. We also discarded the histone deacetylase (HDAC) datasets from consideration, since our focus here was on TFs.

We followed the strategy of Setty and Leslie (Setty and Leslie 2015) to create training and validation data comprising positive and control peaks from each ChIP-Seq dataset. For positive peaks, we took 100 base pair (bp) windows centered at the peak summits. For negative control peaks, we took 100-bp non-peak windows located 100 bp upstream of the positive peaks. Negative control peaks that intersect with positive peaks were discarded along with the corresponding downstream positive peak. For learning motifs, we used the top 2000 positive peaks (ranked by the `signalValue` field) and their associated control peaks, or all positive-control peak pairs if less than 2000 remain. We then randomly shuffled the positive peaks and split them into equal halves (and likewise for control peaks) to obtain our training and validation data.

We used `bwtool` (Pohl and Beato 2014) to extract shape profiles of the positive and control peaks from the bigwig files.

### Identifying promoter-proximal and distal regions

We followed the strategy of Setty and Leslie (Setty and Leslie 2015) to identify promoter-proximal and -distal regions. From UCSC table browser (<u>https://genome.ucsc.edu/cgi-bin/hgTables</u>), we collected the coordinates (hg19) of RefSeq genes (`group=genes` and `gene predictions`, `track=refseq genes`, `table=refgene`, `region=genome`). We then select promoter-proximal regions as the 2-kilobase regions flanking each gene, and the distal regions as the windows spanning 10-kb to 1-Mb regions flanking each gene.

### Sequence motif analysis

We applied the tool `gkm-SVM` (Ghandi, Lee et al. 2014) to compute the sequence motifs that could discriminate between the positive and the control peaks in each training data set. We note that `gkm-SVM` outputs the scores of all 10-mers as predicted by a support vector machine (svm) trained to discriminate between the positive and the control peaks. Thus, `gkm-SVM` does not directly provide a description of a TF’s specificity. To overcome this issue, the `gkm-SVM` package includes an algorithm that iteratively learns a specified number of sequence motifs from the svm scores. We used this algorithm to learn five motifs for each ChIP-seq dataset. Note that, in the original work featuring `gkm-SVM` (Ghandi, Lee et al. 2014), the authors used three motifs to describe specificity of TFs. By utilizing five motifs, we in fact allowed redundancy and presumably weak (low information content) motifs to be included in our analysis in order to ensure that we have a broad sequence-based definition of TF binding sites. We then used the tool `fimo` (Grant, Bailey et al. 2011) to identify all occurrences of these motifs in the positive peaks.

As a second source of sequence-motif hits, we took the genome-wide annotations for occurrences of sequence-motifs curated by Kheradpour and Kellis (Kheradpour and Kellis 2014). In their curated collection, Kheradpour and Kellis used all motifs from available motif libraries and also motifs from several *de novo* motif finders.

### Co-binding regions for a given pair of TFs

For two TFs *f*_1_ and *f*_1_, we first identify the peaks ***S_1_*** of *f*_1_ that intersects with a peak of *f*_2_, and likewise the peaks ***S_2_*** of *f*_2_ that intersects with a peak of *f*_1_. We then merge the genomic regions denoted by ***S_1_*** and ***S_2_*** to obtain the regions where *f*_1_ and *f*_2_ co-bind.

### Calling shape-zingers

A shape-motif ***P*_1_** of a TF *f*_1_ is also a motif for a TF *f*_2_ if ***P*_1_** can discriminate the positive peaks (top 2000 or all if there are less than 2000 positive peaks) of *f*_2_ from control peaks with a hypergeometric *p*-value < 10^−10^. A TF *f*_1_ is a shape-zinger if one or more of its motifs are motifs for at least 30 TFs with a shape-motif in the same cell-line. We chose the fraction 30 following Hunt and Wasserman (Worsley Hunt and Wasserman 2014) who reported sequence-zingers to be enriched in 30-60 datasets.

### Analysis of spacing bias between motif pairs

We followed the strategy of Ng et al. (Ng, Schutte et al. 2014) for analysis of biased spacing between a given pair of motifs. For the given motif pairs, we first compute the distances between their non-overlapping neighboring (adjacent) occurrences. We arbitrarily decide one motif as primary and the other as secondary. A distance between the primary-secondary motif pair is positive if the primary-motif occurs upstream of the secondary-motif, zero if they occur at the same location, and it is negative otherwise. We test each distance between -50 to +50 bps, and call a distance to be significant if the binomial *p*-value (after Bonferroni correction) is significant (< 10^−5^).

### ShapeAmotif analysis for TF families

We took the classification of TFs into families according to their DNA-binding domains from `TFClass` (Wingender, Schoeps et al. 2013). For each TF in a family, we first selected the sets of peaks that do not overlap with the peaks of any other TF in the same family. We then used ShapeMF on the remaining sets of peaks to identify the shape motifs of each TF.

## Acknowledgement

NIH/NHLBI Bench to Bassinet Program UM1HL098179, to K.S.P. and B.G.B, and by William H. Younger, Jr. (B.G.B.)

## Supplementary Figure Legends

**Supplementary Figure 1.**
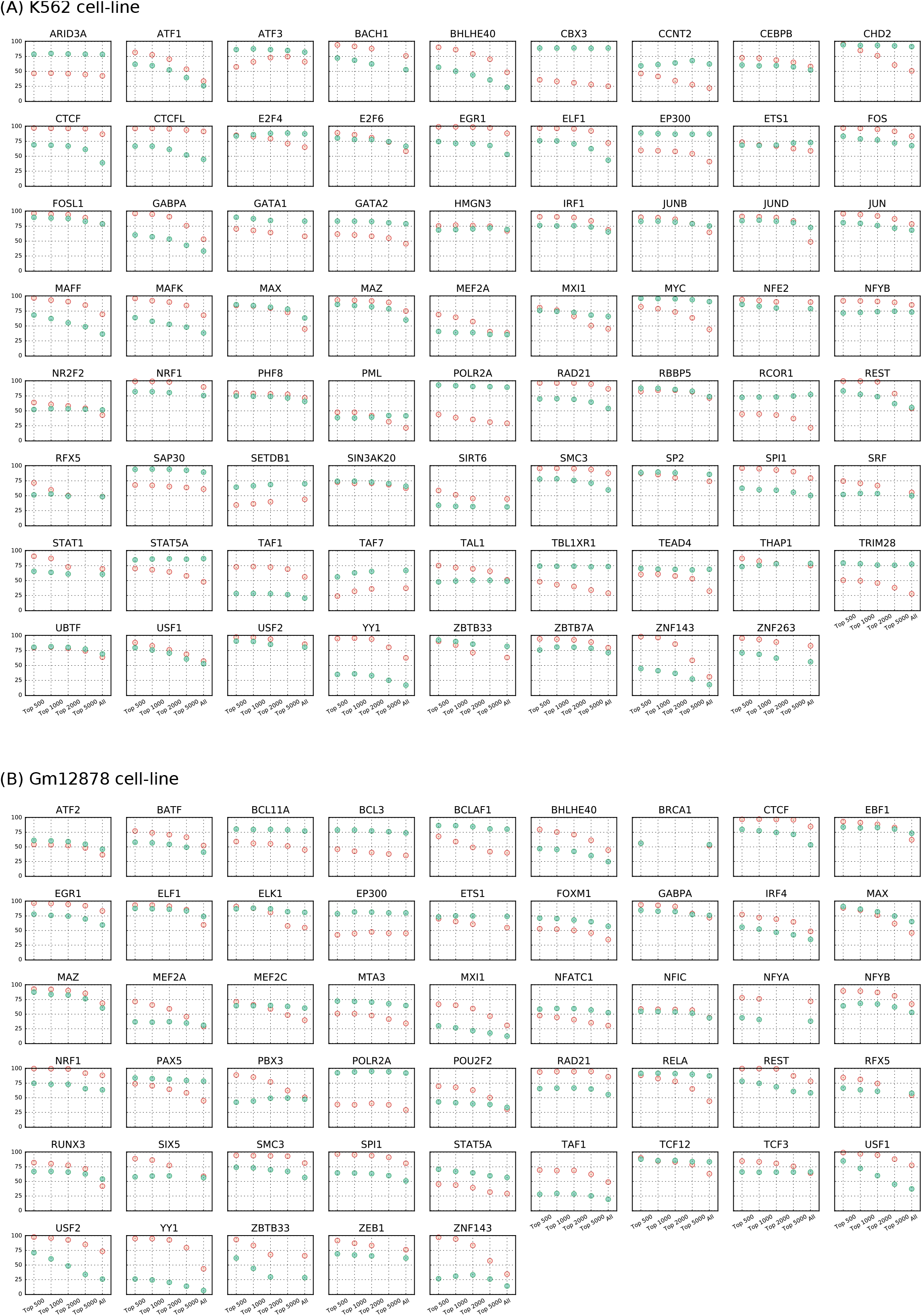
Fractions of top 500, 1000, 2000, 5000, and all peaks of a TF’s ChIP-Seq dataset that contain an occurrence of the TF’s sequence-motif (orange circle) and shape-motif (green-circle). Data plotted for TFs assayed in (A) K562 and (B) Gm12878 cell-lines.

**Supplementary Figure 2.**
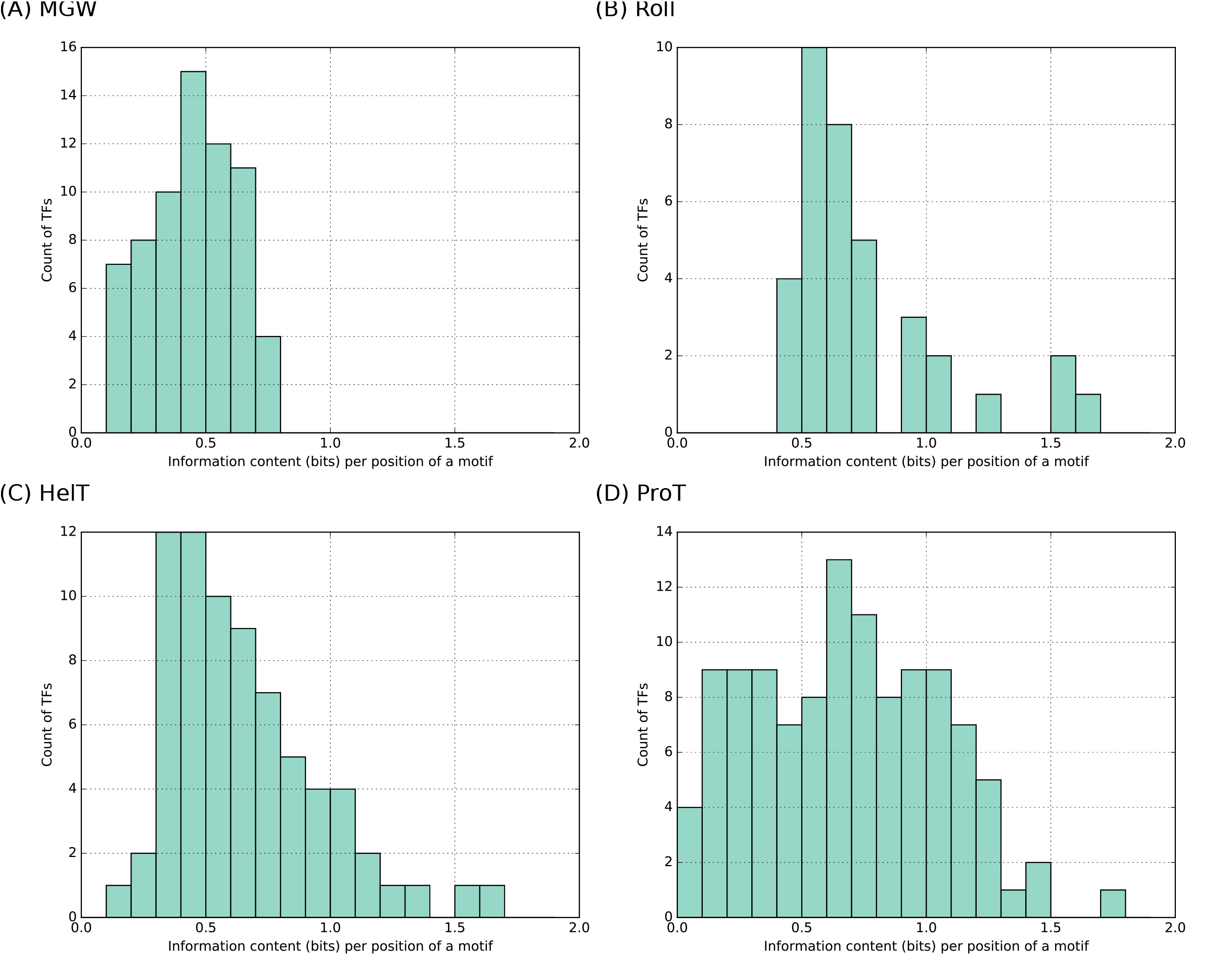
Histograms showing number of motifs of the four different features having different values of Information Content per Position (ICP). The ICP statistic for a shape-motif is derived from the sequences underlying the occurrences of the shape-motif (Methods).

**Supplementary Figure 3.**
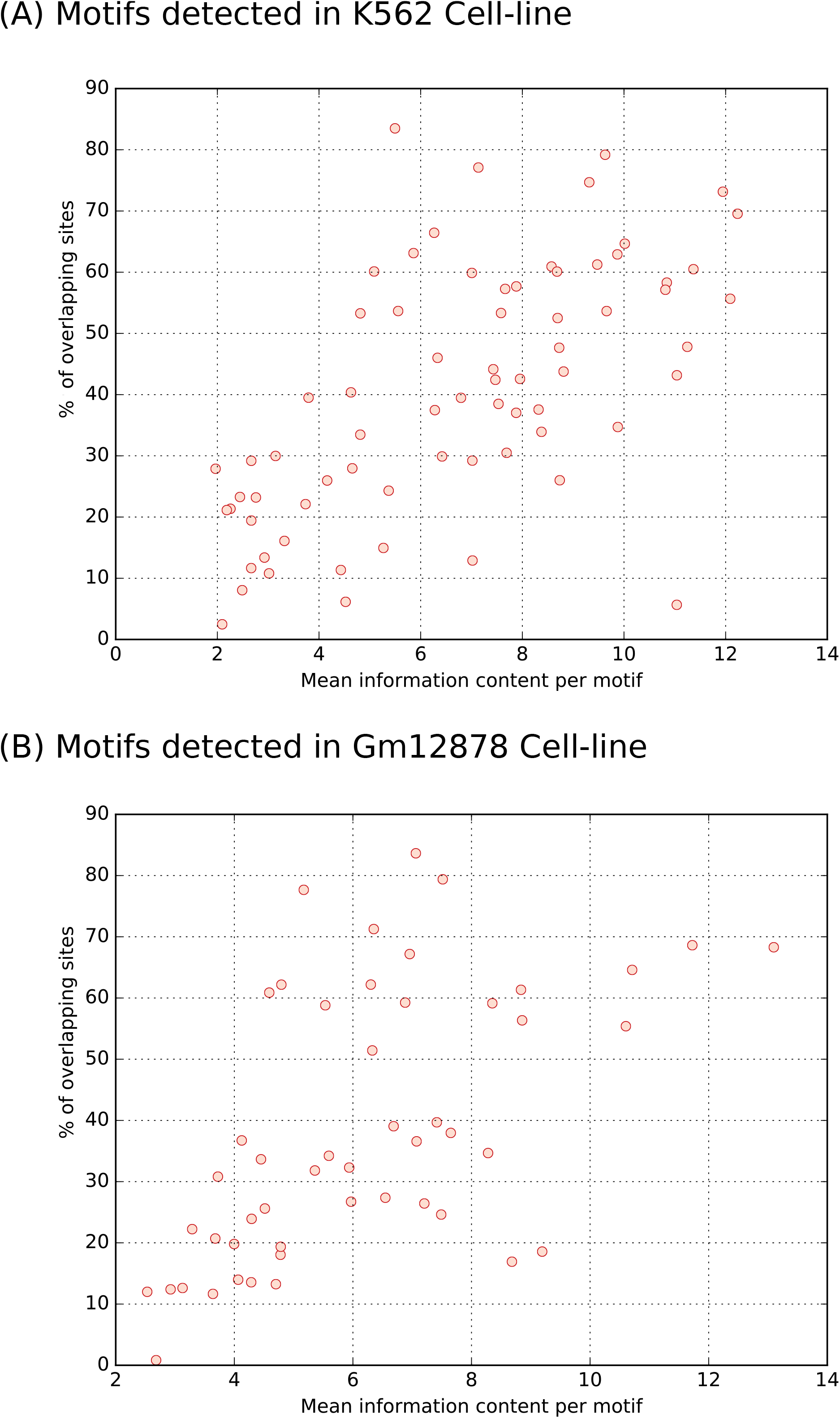
Scatterplot showing a monotonically increasing relationship between the fraction of overlapping sites of a TF and the mean information content derived from sequence-logos underlying its shape-motifs. Each circle denotes a TF and the mean information content is the mean of the information contents of its different shape-motifs. Data shown for TFs in (A) K562 and (B) Gm12878 cell-lines.

**Supplementary Figure 4.**
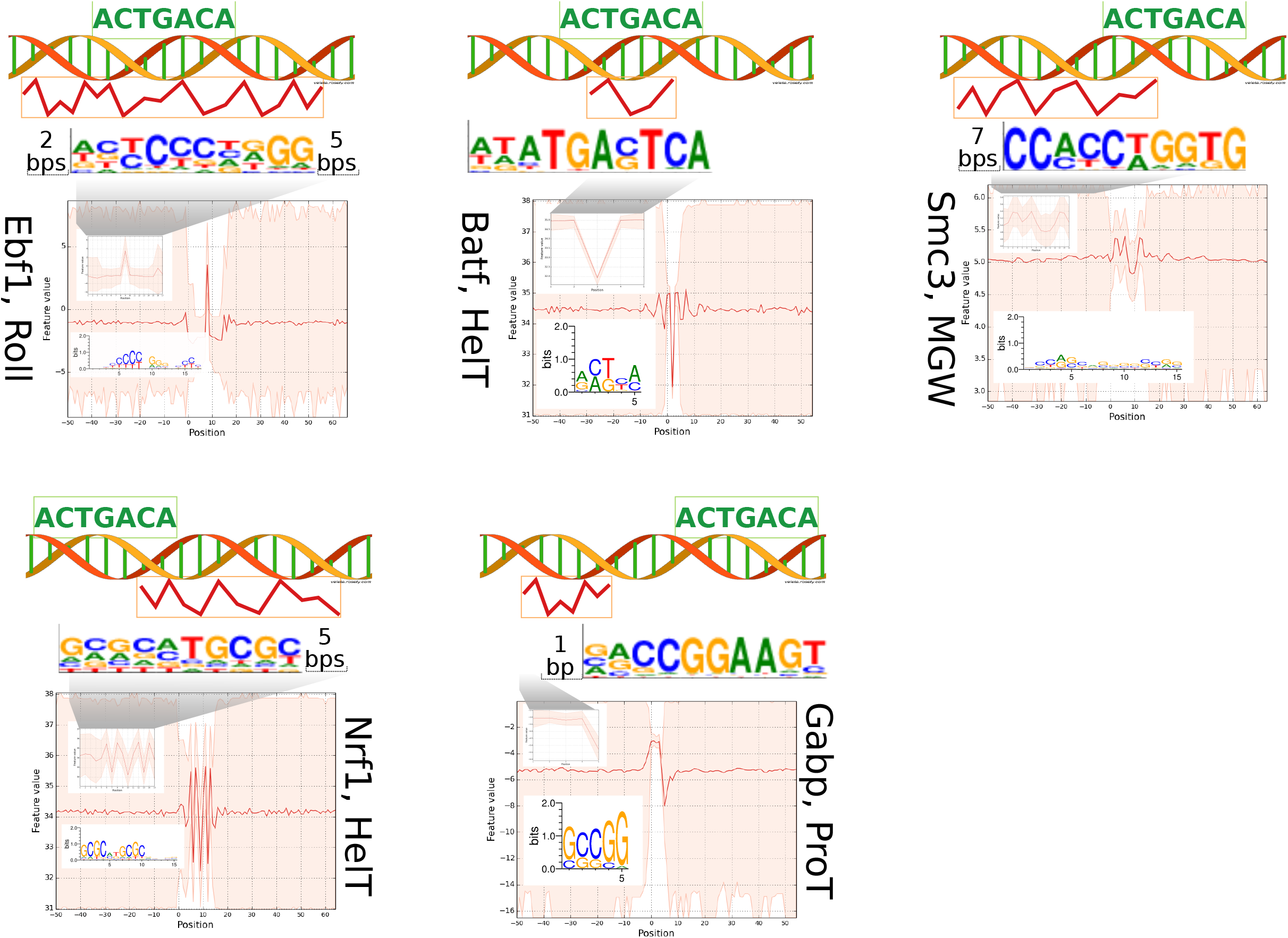
Different scenarios of shape- and sequence-motif co-occurrence found enriched in datasets from the Gm12878 cell-line.

**Supplementary Figure 5.**
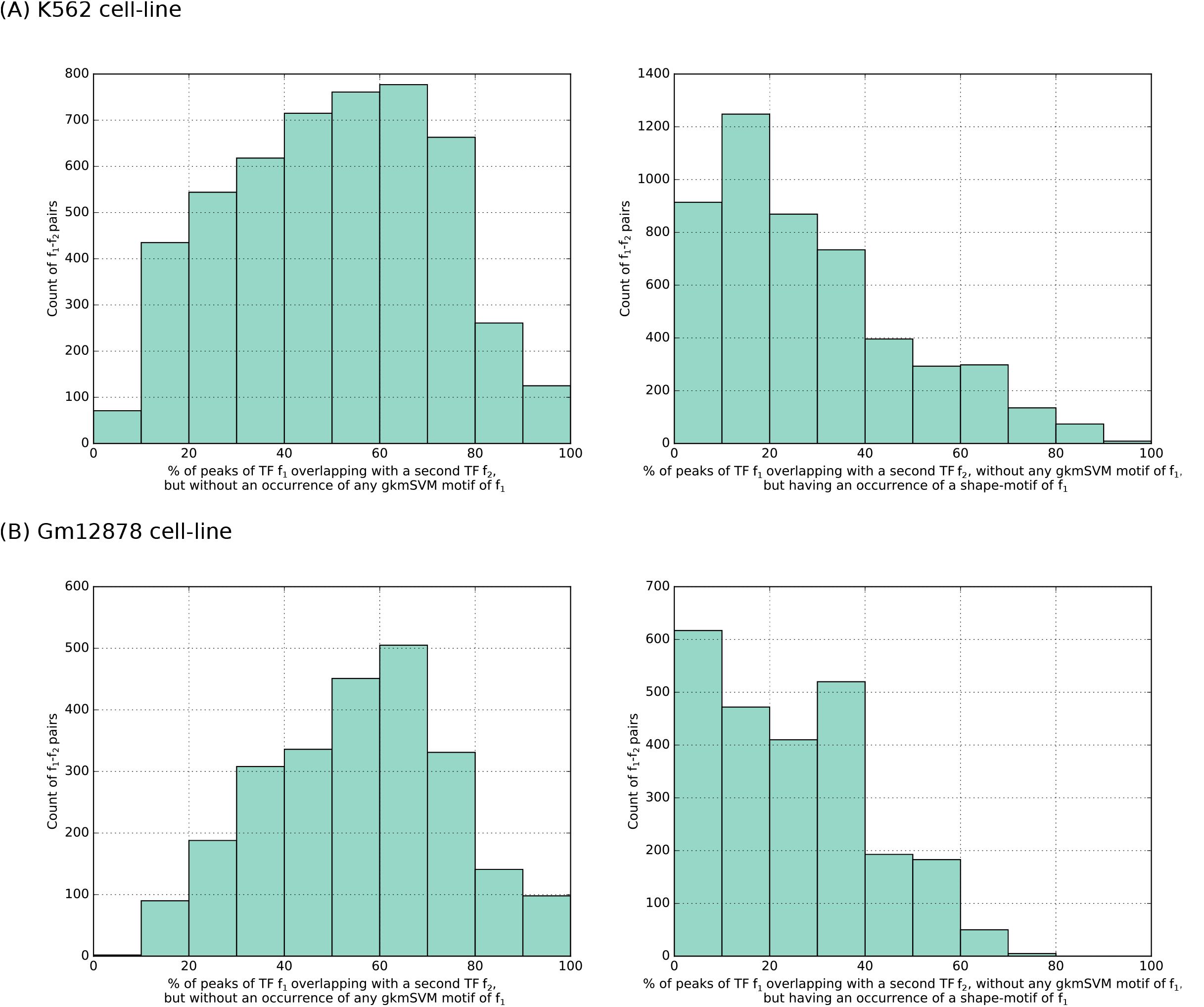
Histograms showing number of co-binding TF-pairs (f1, f2) for different fractions of peaks of f1 lacking a sequence-motif of f1 (left panel) or containing a shape-motif of f1 (right panel) in (A) K562 and (B) Gm12878 cell-lines.

**Supplementary Figure 6.**
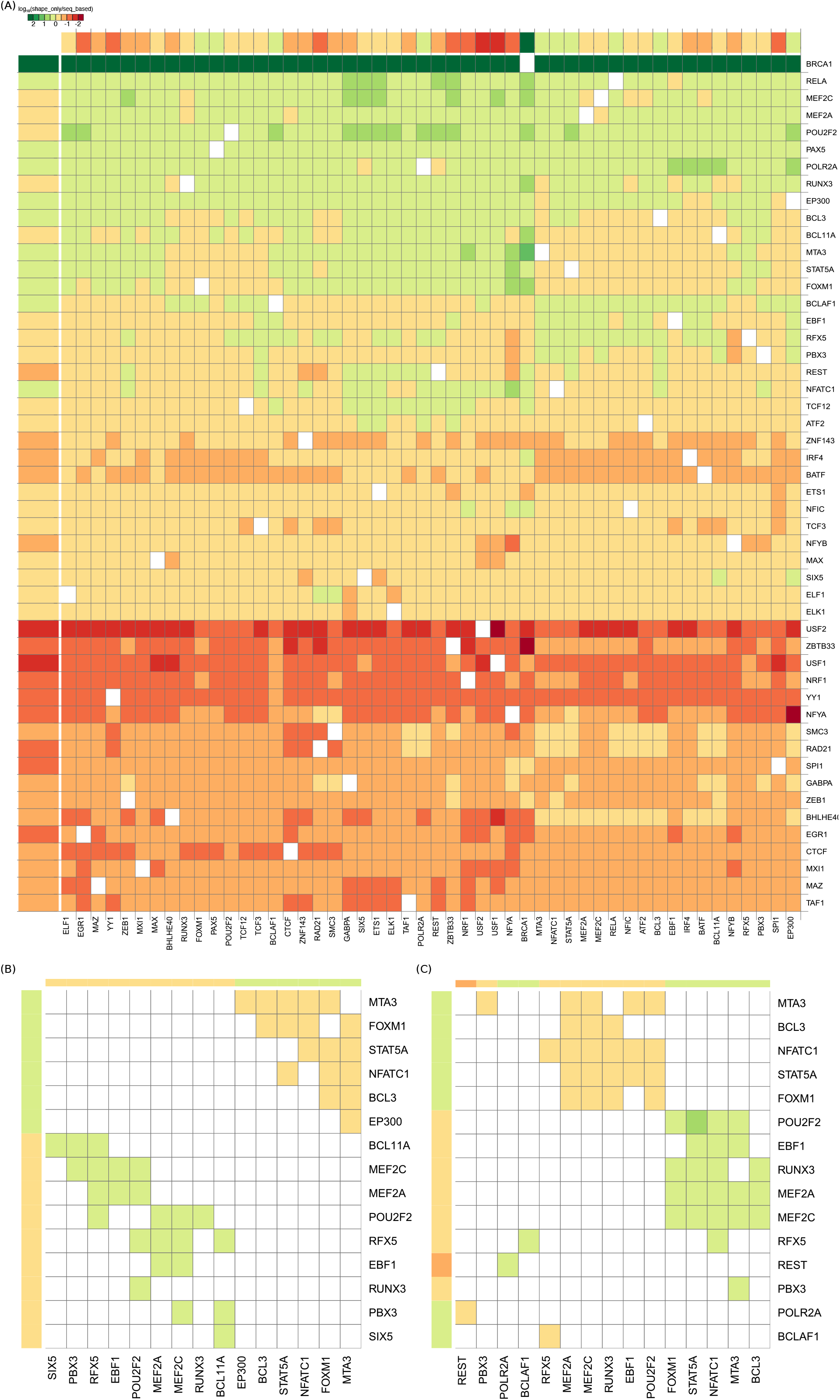
Heatmaps similar to Figure 5 showing results of our co-binding analysis on TF ChIP-Seq data in Gm12878 cell-line. (A) Co-binding TF pairs often utilizing shape-specific binding. The binding mode of a TF in a co-binding region may alter depending on its partner: both (B) or one (C) TF may alter binding mode.

**Supplementary Figure 7.**
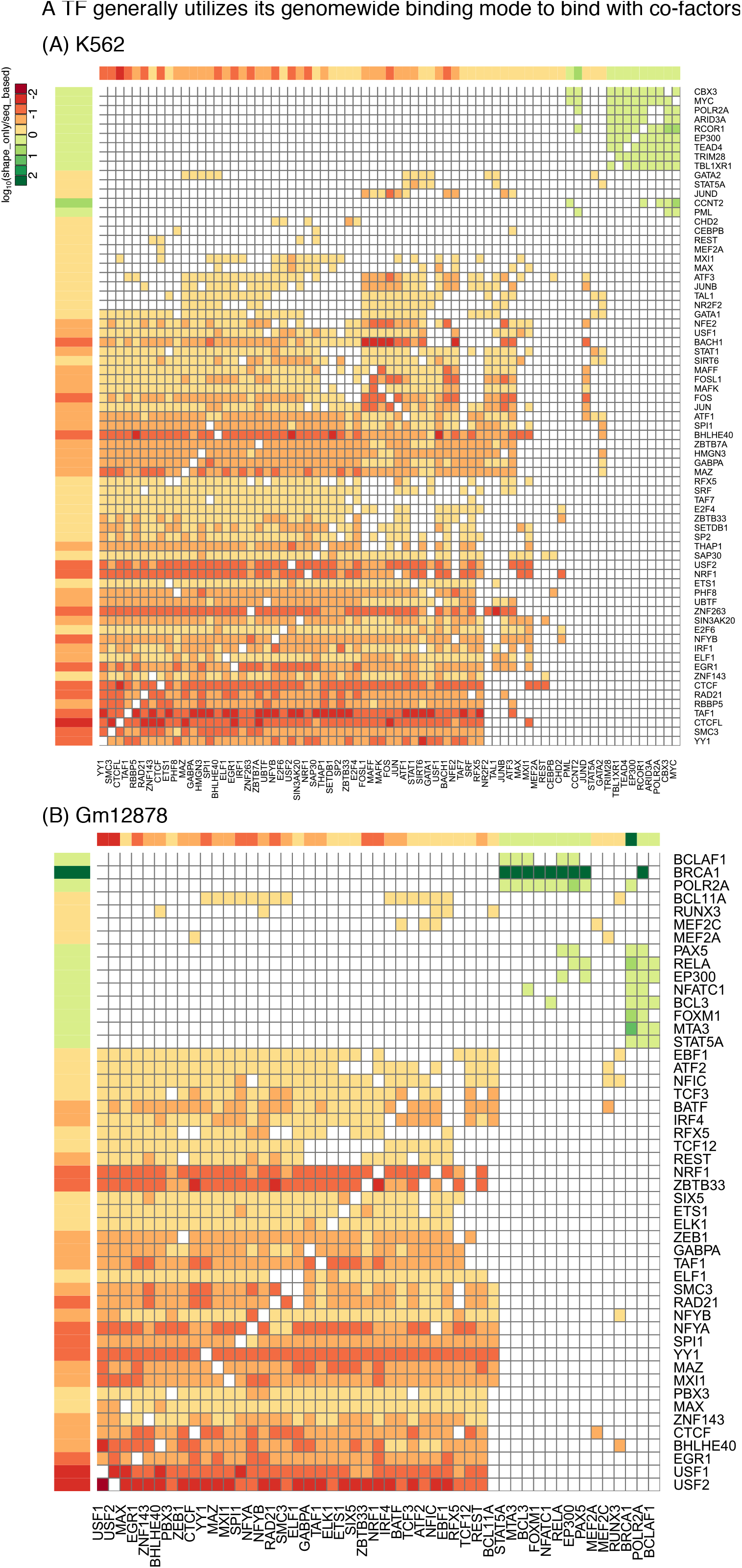
Heatmaps similar to Figure 5 showing that a TF generally maintains its genomewide binding mode when co-binding with other TFs in (A) K562 and (B) Gm12878 cell-lines.

**Supplementary Figure 8.**
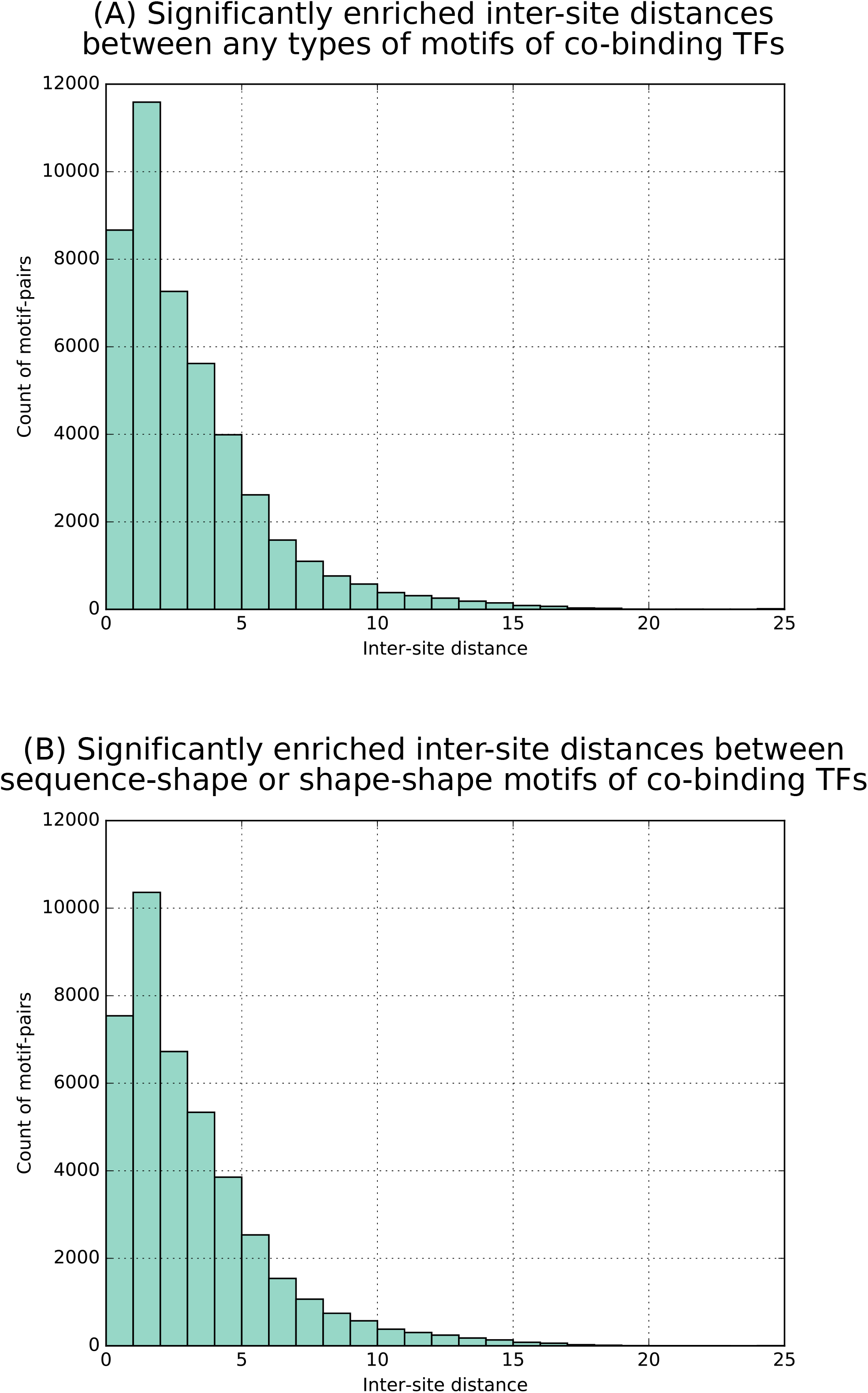
Significantly enriched inter-site distances between (A) any types of motif-pairs and (B) shape-sequence or shape-shape motif-pairs. Instances from K562 and Gm12878 cell-lines were combined in each panel.

**Supplementary Figure 9.**
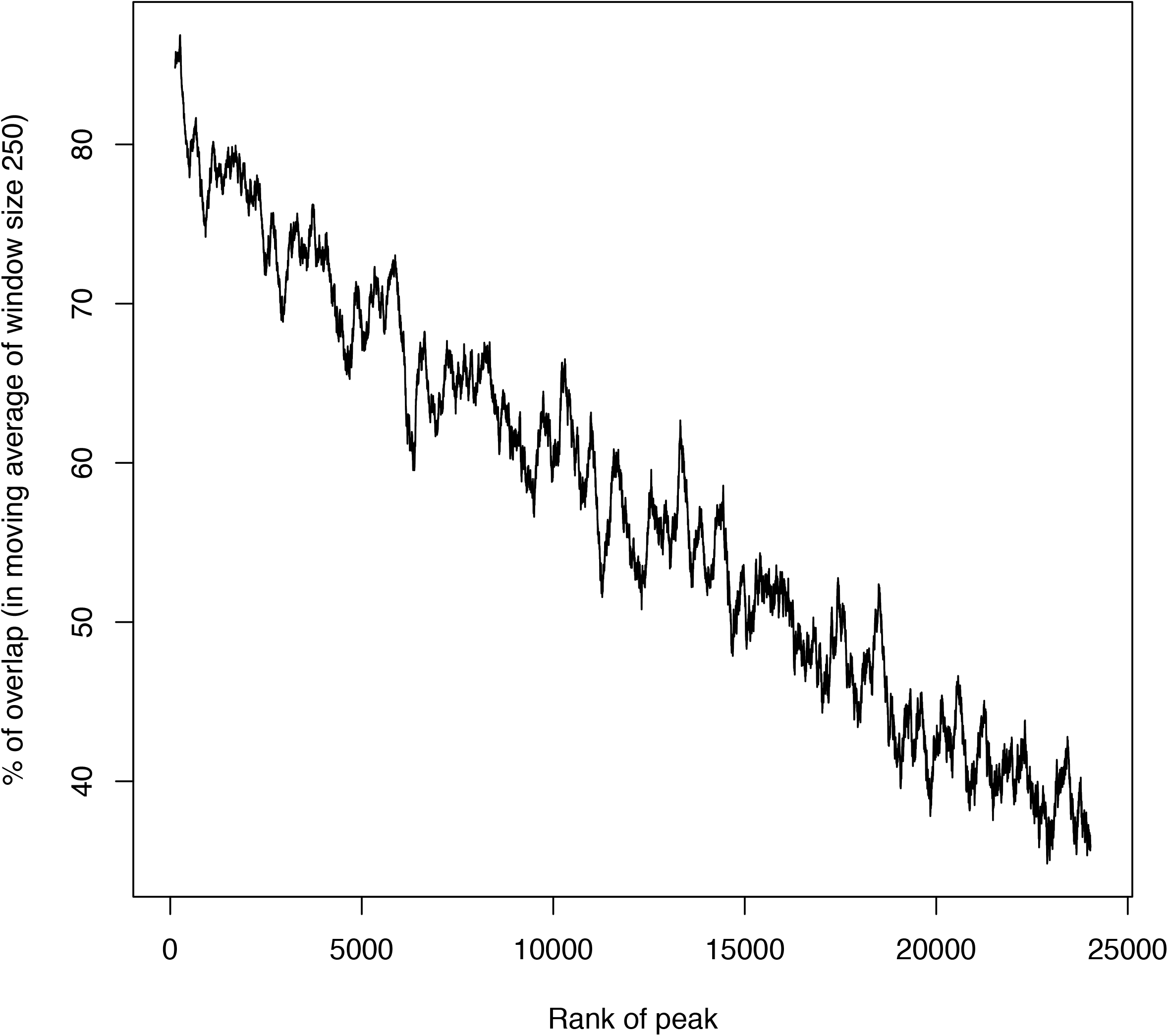
Moving average plot (window size = 250) showing monotonically decreasing relationship between rank of MYC peaks and their of overlap (bps) with MAX peaks.

**Supplementary Figure 10.**
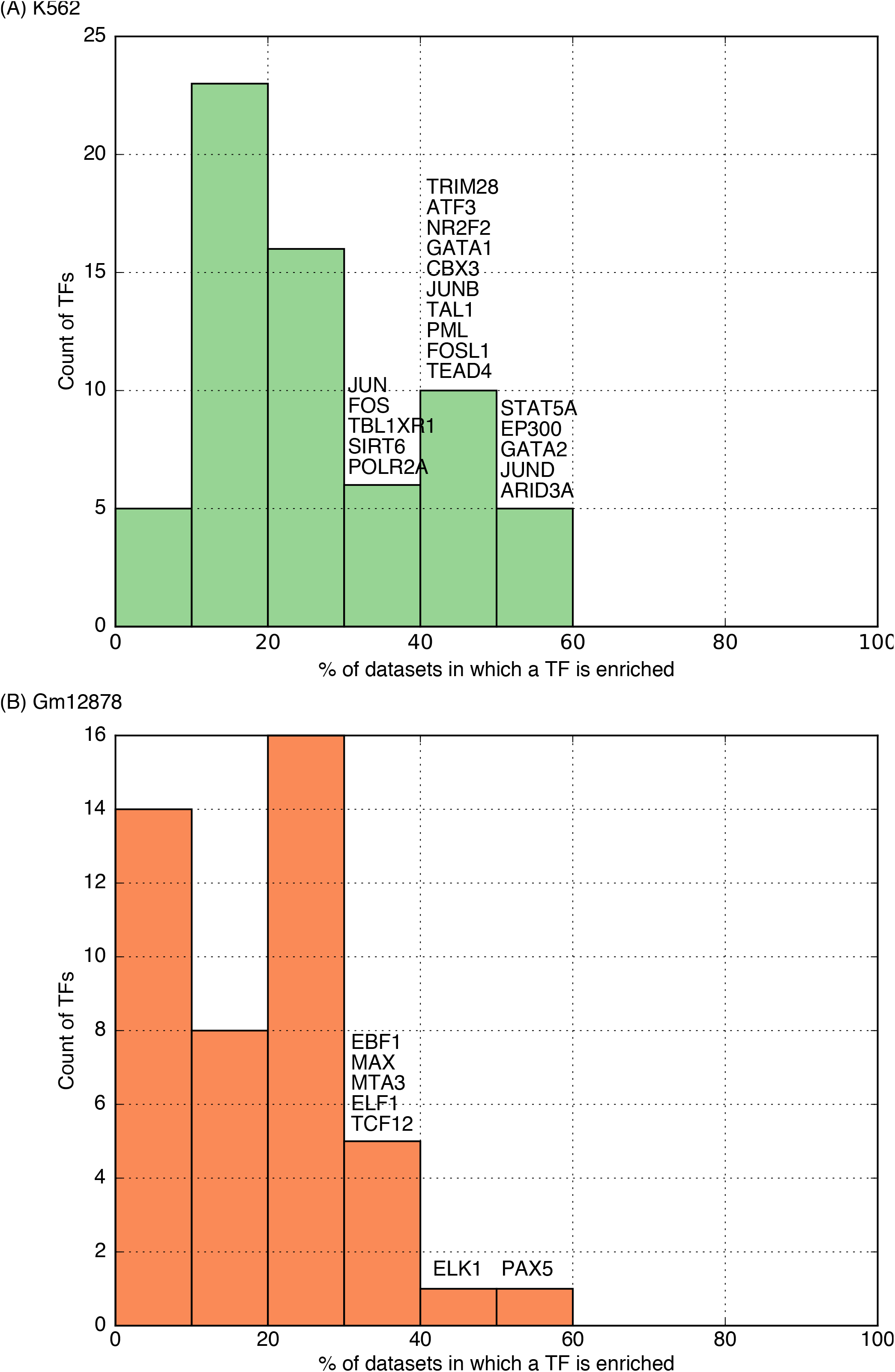
Histograms of the numbers of TFs whose shape-motifs are enriched in different fractions of ChIP-Seq datasets in (A) K562 and (B) Gm12878 cell-lines. Shape-zingers are defined as the TFs with enrichment in at least 30 datasets (Methods).

**Supplementary Figure 11.**
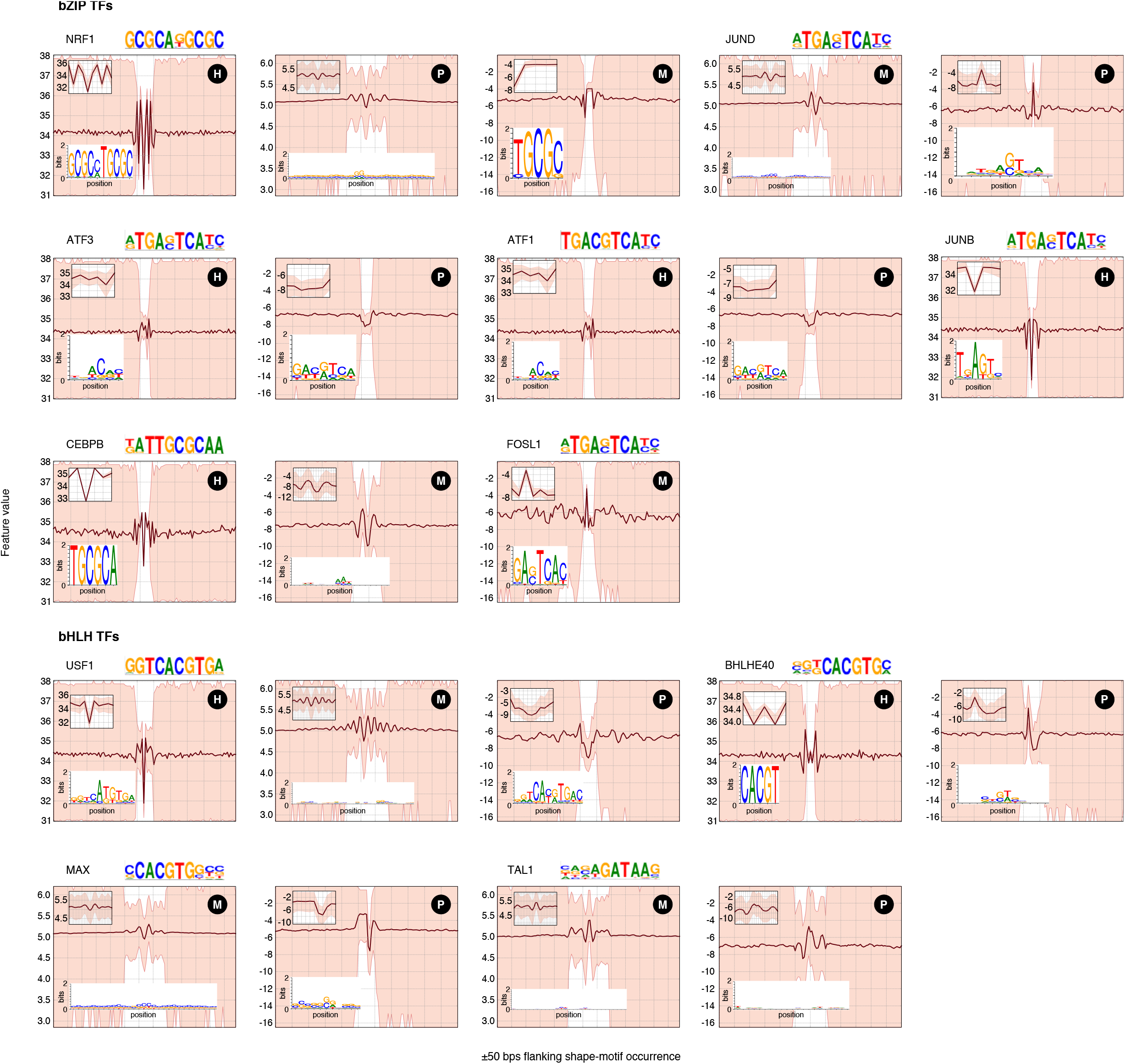
Shape-motifs of bZIP and bHLH proteins (for which ChIP-Seq assays were performed by ENCODE in the K562 cell-line and we could find a shape-motif) showing that TFs within the same family extensively utilize different shape-motifs (H, M, and P denote motifs for HelT, MGW, and ProT, respectively), and/or combinations of different shape-features to recognize their target binding sites. A feature is not mentioned for a TF if the TF does not have a shape-motif for that feature. Seven of 13 bZIP TFs and 4 of 7 bHLH TFs were found to have a shape-motif in this analysis.

**Supplementary Figure 12.**
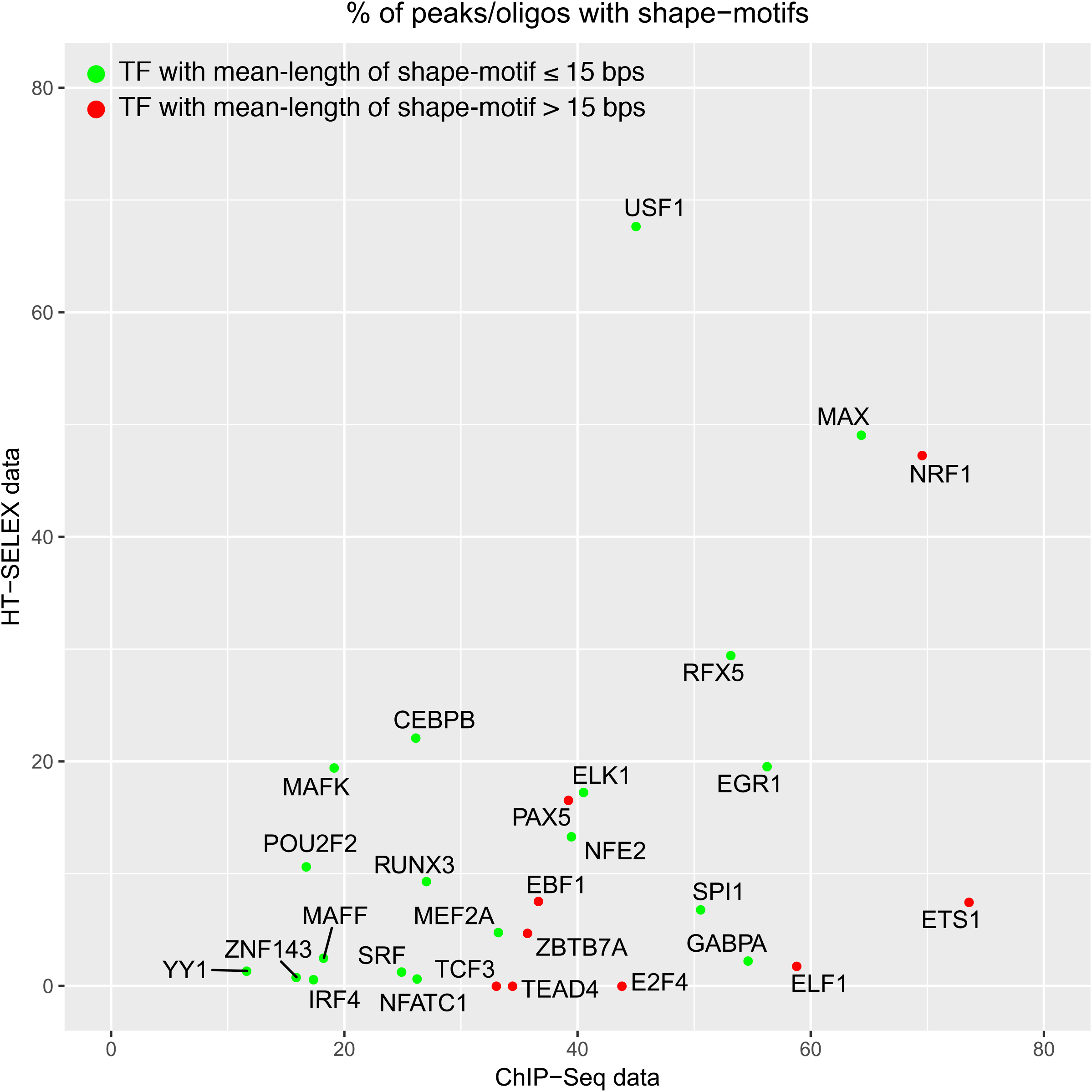
Scatterplot showing fraction of ChIP-peaks of a TF with its shape-motifs vs. fraction of the TF’s HT-SELEX oligonucleotides with its shape-motifs. Green/red data points indicate TFs with mean-length of shape-motifs smaller/longer than 15 bps.

**Supplementary Table 1:**
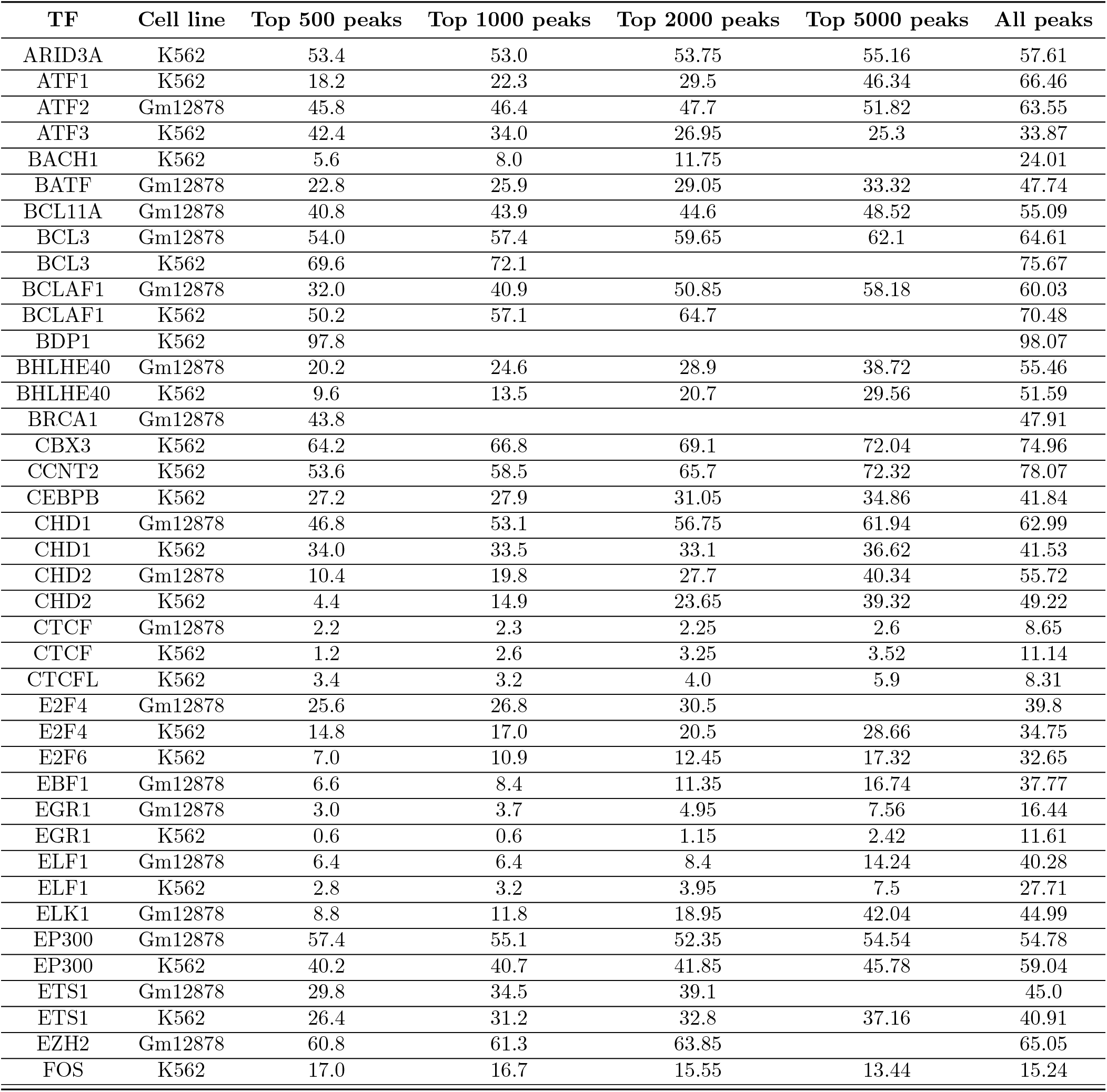
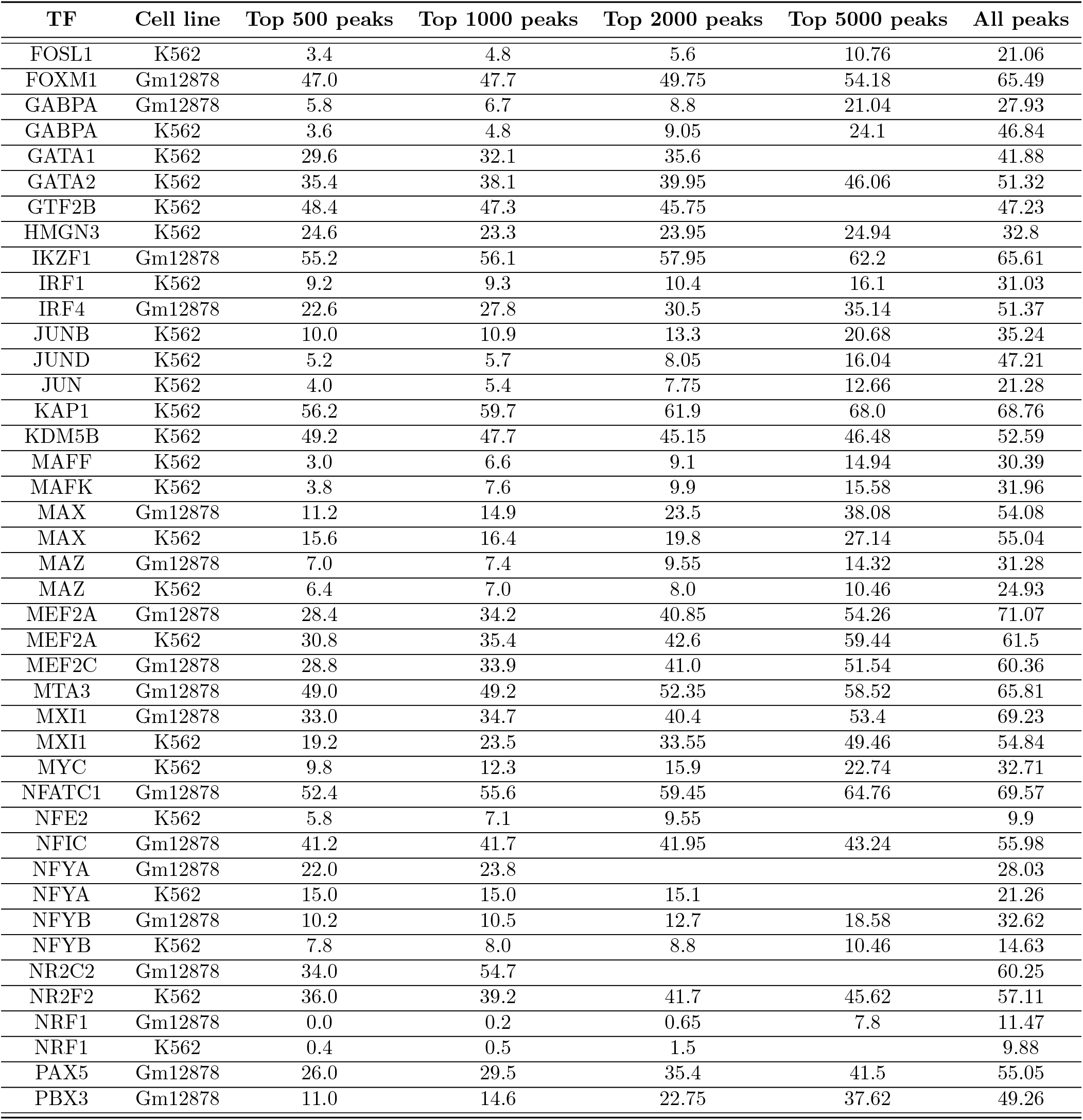
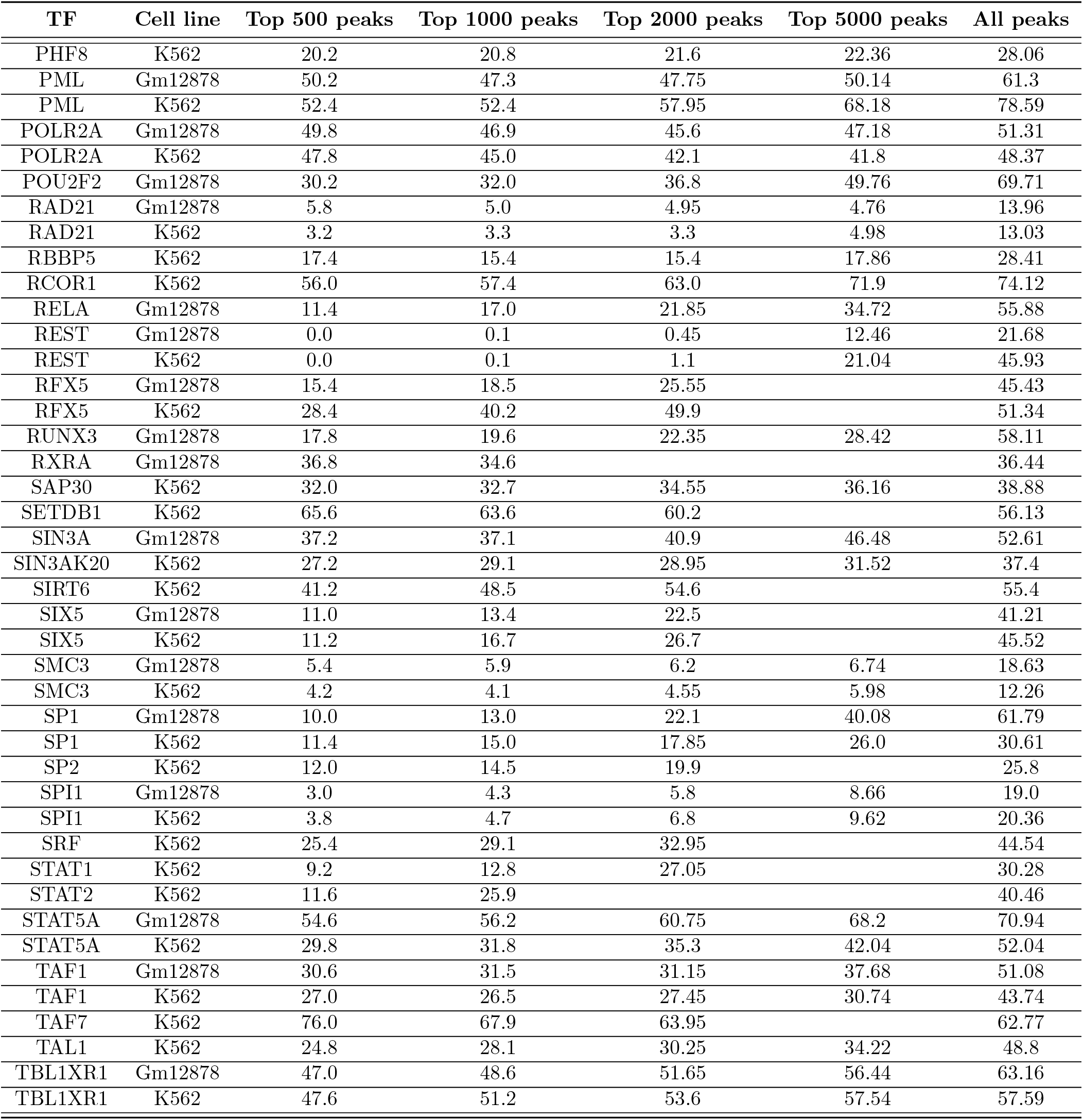
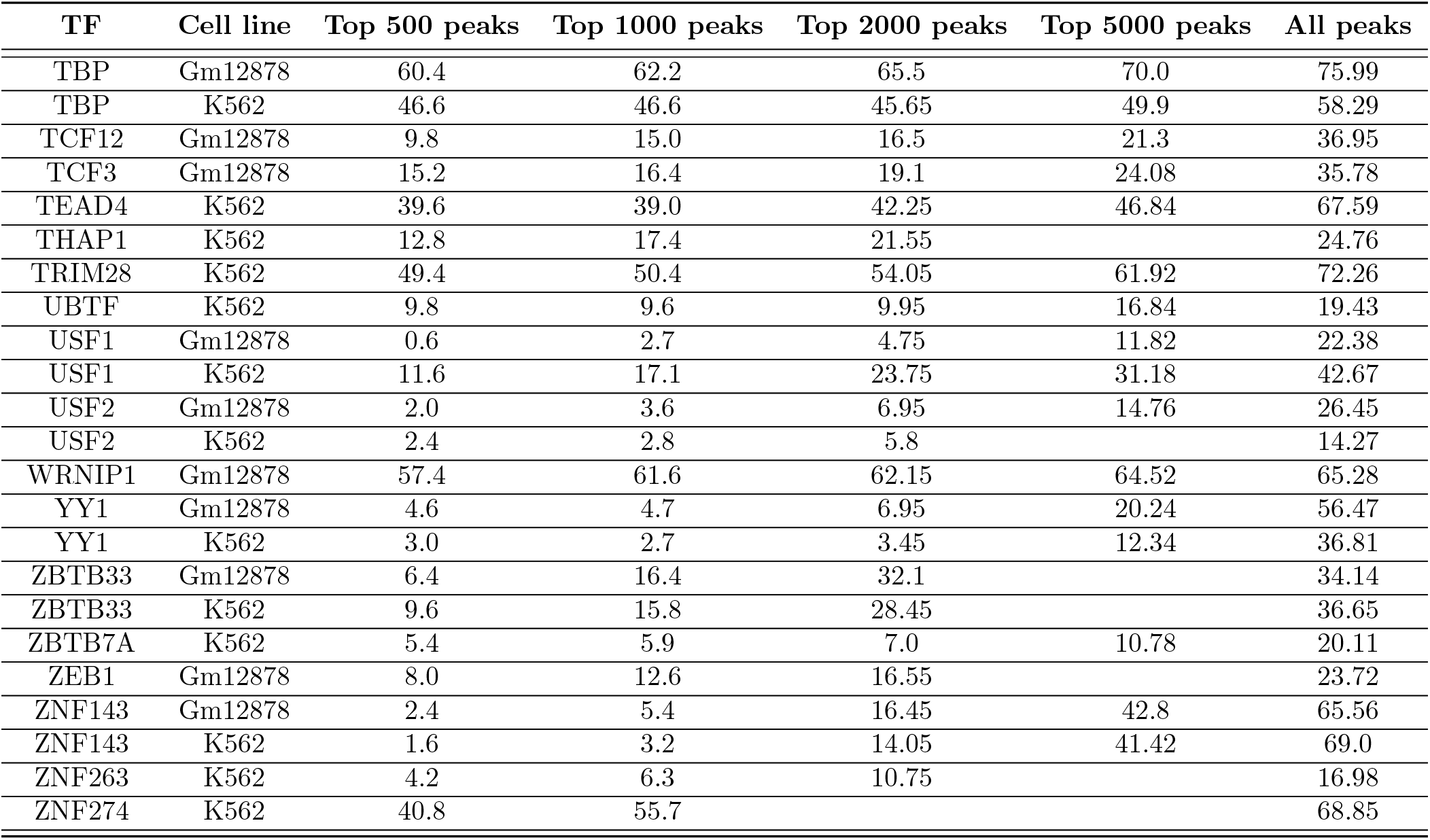
Fraction (%) of peaks without occurrence of any sequence motif

**Supplementary Table 2:**
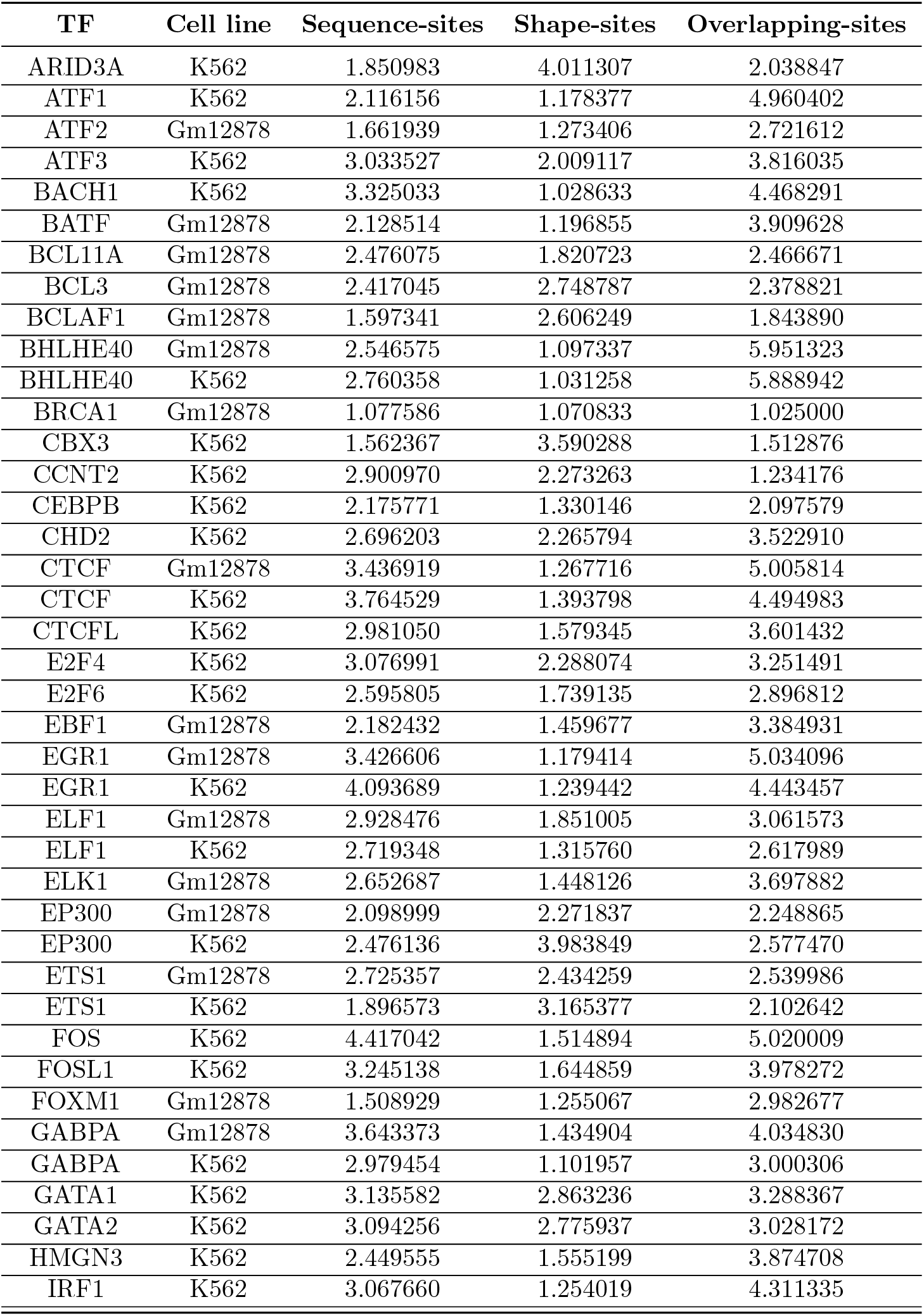
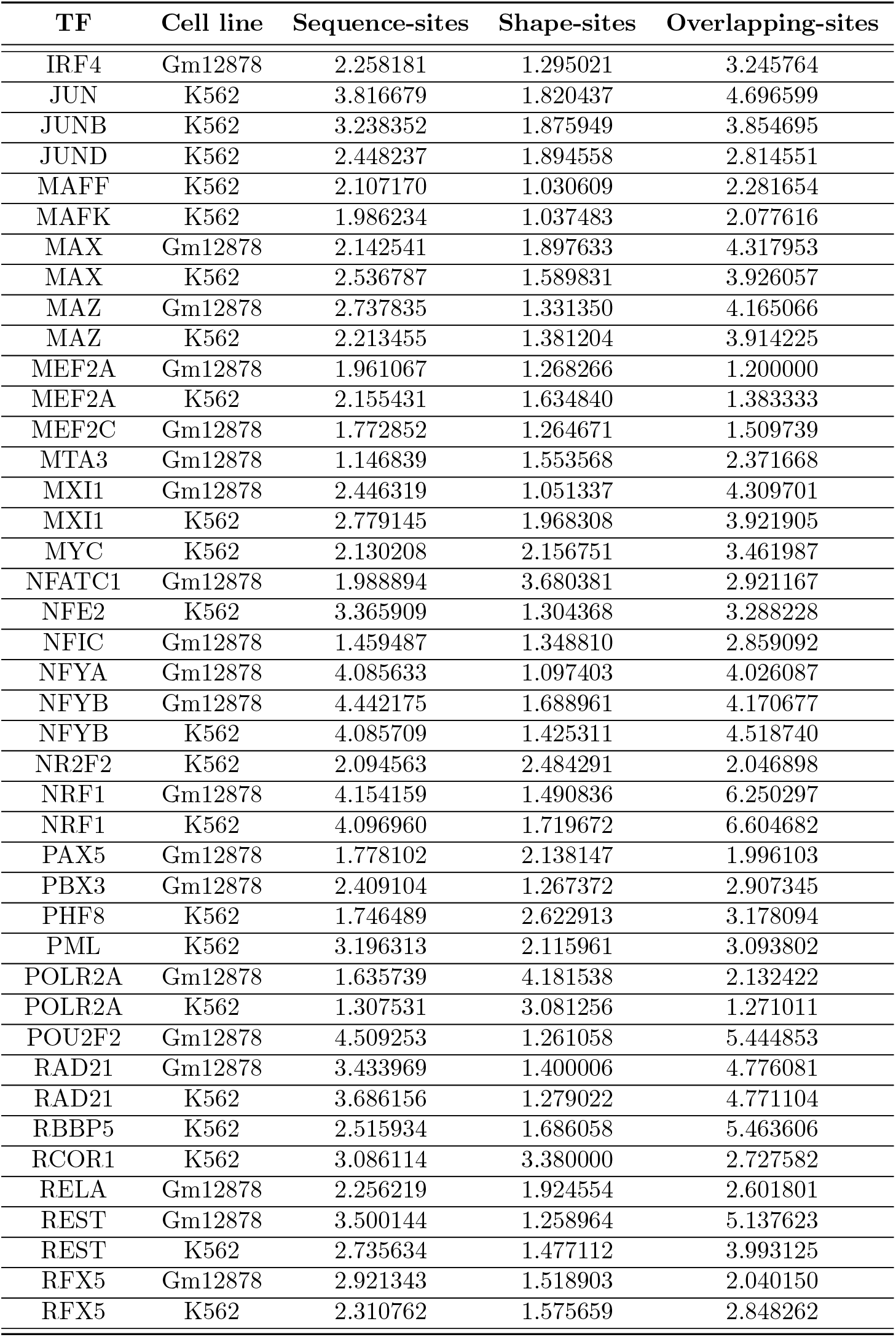
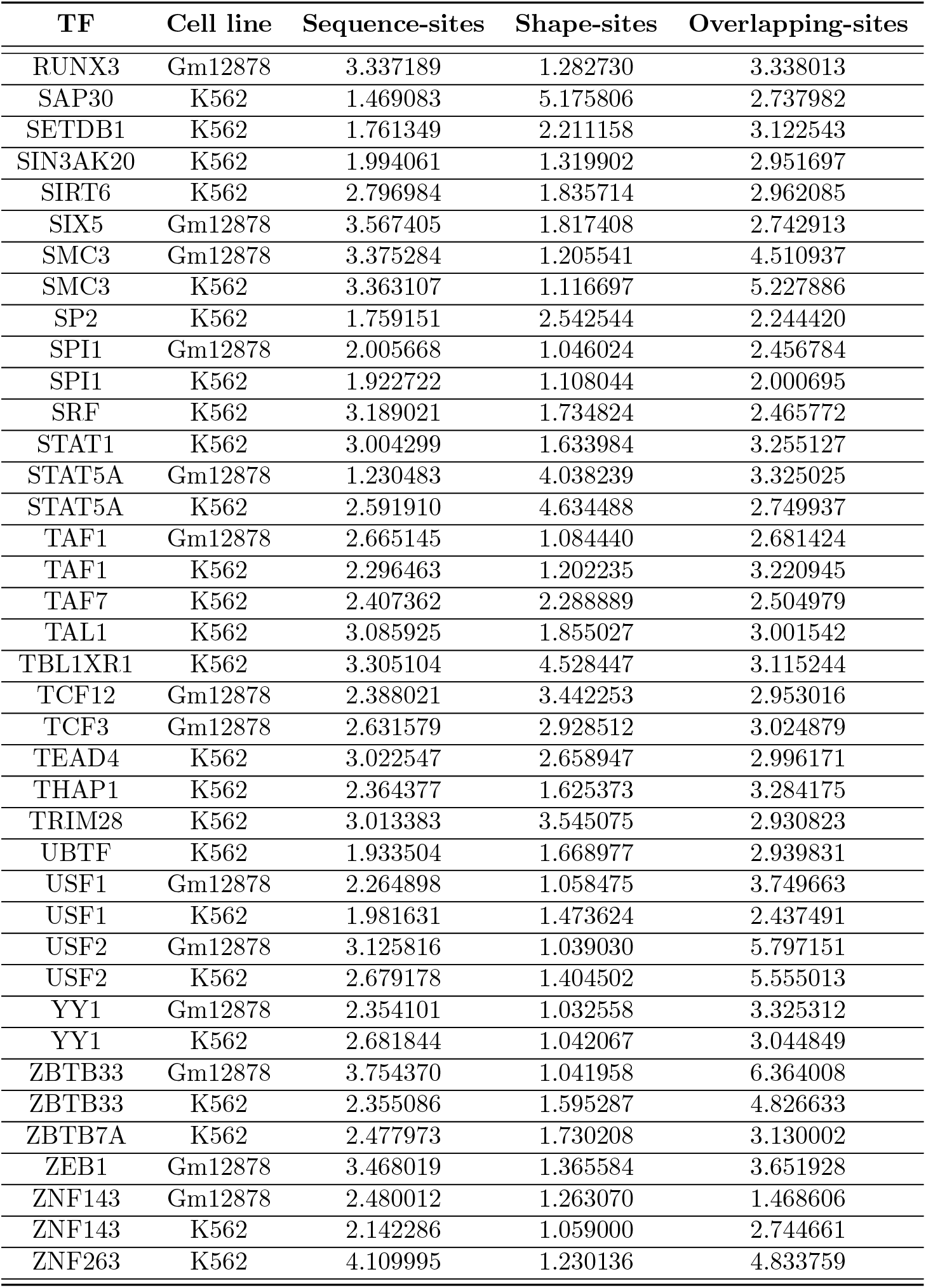
Average number of sequence-, shape-, and overlapping-sites per peak

**Supplementary Table 3:** Information content (bits) for each type of motif of each TF

| TF | Cell line | HelT motif | MGW motif | ProT motif | Roll motif |
| --- | --- | --- | --- | --- | --- |
| ATF1 | K562 | 4.354304 |  | 5.813853 |  |
| ATF2 | Gm12878 | 3.739719 |  | 3.706825 |  |
| ATF3 | K562 | 3.686794 | 5.597207 | 4.679672 |  |
| BACH1 | K562 | 8.401840 | 8.373654 | 4.629956 |  |
| BATF | Gm12878 | 4.755129 |  | 4.831843 |  |
| BCL11A | Gm12878 | 2.731374 |  | 2.331119 |  |
| BCL3 | Gm12878 | 8.307989 | 4.415979 | 8.087002 | 13.915730 |
| BCLAF1 | Gm12878 |  | 6.153377 | 5.784666 |  |
| BHLHE40 | Gm12878 | 6.659689 |  | 6.043686 |  |
| BHLHE40 | K562 | 4.831120 |  | 6.158475 |  |
| BRCA1 | Gm12878 | 7.902666 |  | 13.303756 |  |
| CBX3 | K562 |  | 3.974170 | 2.058881 |  |
| CCNT2 | K562 |  | 13.223148 | 11.681911 | 8.210620 |
| CEBPB | K562 | 3.033718 |  | 2.300033 |  |
| CHD2 | K562 | 5.157121 | 8.678947 | 7.214341 |  |
| CTCF | Gm12878 |  | 9.591501 | 6.342796 | 10.569503 |
| CTCF | K562 |  | 8.657654 | 8.240361 | 11.524574 |
| CTCFL | K562 |  | 11.342471 | 14.267542 | 11.094728 |
| E2F4 | K562 | 7.629851 | 12.680834 | 12.811067 |  |
| E2F6 | K562 | 8.058094 | 10.192821 | 11.383686 |  |
| EBF1 | Gm12878 | 7.546488 | 8.776658 | 3.714710 | 8.257570 |
| EGR1 | Gm12878 | 8.707555 | 14.324367 | 12.780700 | 11.083092 |
| EGR1 | K562 | 8.566762 | 14.611101 | 10.949566 | 10.856076 |
| ELF1 | Gm12878 | 5.410424 |  | 8.677957 | 7.525383 |
| ELF1 | K562 | 6.040370 |  | 9.723409 | 9.366509 |
| ELK1 | Gm12878 | 5.746806 |  | 6.420904 | 7.473470 |
| EP300 | Gm12878 | 3.162812 |  | 2.687594 |  |
| EP300 | K562 |  | 3.287761 | 2.046333 |  |
| ETS1 | Gm12878 |  | 8.301907 | 6.676335 |  |
| ETS1 | K562 |  | 10.002779 | 7.620823 |  |
| FOS | K562 | 5.182979 | 4.617328 | 7.777137 |  |
| FOSL1 | K562 | 5.021982 | 5.388716 | 8.386084 |  |
| FOXM1 | Gm12878 | 4.213639 |  | 3.916519 |  |
| GABPA | Gm12878 | 5.636910 |  | 8.110776 | 9.203928 |
| GABPA | K562 | 5.335118 |  | 8.716376 | 8.357448 |
| GATA1 | K562 |  | 4.458784 | 3.010830 |  |
| GATA2 | K562 |  | 3.702154 | 2.937098 |  |
| HMGN3 | K562 |  | 14.511307 | 12.863219 | 8.898824 |
| IRF1 | K562 | 5.878582 | 7.743200 |  | 12.554550 |
| IRF4 | Gm12878 | 4.088219 |  | 4.161863 |  |
| JUN | K562 | 3.888448 | 5.250780 | 5.300358 |  |
| JUNB | K562 | 5.128315 | 3.345063 | 5.411392 |  |
| JUND | K562 | 4.050870 | 4.431581 | 5.952647 |  |
| MAFF | K562 | 6.562588 | 6.107544 | 8.285775 | 7.070965 |
| MAFK | K562 | 6.510408 | 7.551979 | 9.655213 | 10.579278 |
| MAX | Gm12878 | 5.598401 | 8.576249 | 8.068755 |  |
| MAX | K562 | 5.315556 | 8.767118 | 8.989175 |  |
| MAZ | Gm12878 | 7.873606 | 11.101825 | 13.699716 | 10.163128 |
| MAZ | K562 | 7.908911 | 12.827615 | 12.629814 | 10.002484 |
| MEF2A | Gm12878 | 2.680824 |  |  |  |
| MEF2A | K562 |  |  | 2.094583 |  |
| MEF2C | Gm12878 | 4.263320 |  | 4.305974 |  |
| MTA3 | Gm12878 | 2.676849 |  | 4.681152 |  |
| MXI1 | Gm12878 | 5.534359 |  |  |  |
| MXI1 | K562 | 4.396907 | 10.588809 | 4.277373 |  |
| MYC | K562 | 6.051838 | 9.564069 | 7.372048 | 5.091212 |
| NFATC1 | Gm12878 | 3.754768 |  | 5.802839 |  |
| NFE2 | K562 | 3.061157 | 4.814266 | 3.505833 |  |
| NFIC | Gm12878 | 3.927508 |  | 5.102922 |  |
| NFYA | Gm12878 | 4.589229 |  |  |  |
| NFYB | Gm12878 |  | 5.594612 |  |  |
| NFYB | K562 | 3.359020 | 7.754133 |  |  |
| NR2F2 | K562 |  |  | 2.257454 |  |
| NRF1 | Gm12878 | 11.949359 | 13.625259 | 13.717602 |  |
| NRF1 | K562 | 10.288648 | 11.391690 | 12.420220 |  |
| PAX5 | Gm12878 | 3.373766 | 5.922113 | 3.578923 |  |
| PBX3 | Gm12878 | 4.540342 |  | 4.359496 |  |
| PHF8 | K562 |  |  | 11.945955 |  |
| PML | K562 |  |  | 2.488596 |  |
| POLR2A | Gm12878 |  |  | 4.701124 |  |
| POLR2A | K562 |  | 6.291600 | 4.655549 | 4.853058 |
| POU2F2 | Gm12878 | 3.639135 |  |  |  |
| RAD21 | Gm12878 |  | 5.435081 | 3.938752 | 9.532026 |
| RAD21 | K562 |  | 8.654341 | 6.573681 | 10.820522 |
| RBBP5 | K562 |  | 11.892422 | 8.139989 |  |
| RCOR1 | K562 |  | 5.947349 | 3.099629 |  |
| RELA | Gm12878 | 2.811867 |  |  | 6.753642 |
| REST | Gm12878 | 7.511681 | 7.408559 | 8.465631 | 10.018286 |
| REST | K562 | 6.879823 | 8.100623 | 8.063087 | 8.775688 |
| RFX5 | Gm12878 | 2.541426 |  | 3.712873 |  |
| RFX5 | K562 | 2.805591 |  | 3.482983 |  |
| RUNX3 | Gm12878 | 3.491183 |  | 3.087217 |  |
| SAP30 | K562 |  |  | 7.878489 |  |
| SETDB1 | K562 |  | 8.650117 | 6.412054 |  |
| SIN3AK20 | K562 | 5.404872 | 8.604263 | 8.733767 |  |
| SIRT6 | K562 |  |  | 1.964513 |  |
| SIX5 | Gm12878 | 5.358556 |  |  |  |
| SMC3 | Gm12878 | 4.986087 | 7.022509 | 5.397144 | 10.118401 |
| SMC3 | K562 | 5.406092 | 9.008039 | 6.745234 | 10.366183 |
| SP2 | K562 |  | 7.612506 | 9.026121 |  |
| SPI1 | Gm12878 | 6.526468 |  | 7.895665 | 6.451772 |
| SPI1 | K562 | 7.134858 |  | 8.053192 | 7.799134 |
| SRF | K562 |  |  | 4.160145 |  |
| STAT1 | K562 |  |  | 4.624593 | 7.933305 |
| STAT5A | Gm12878 |  |  | 3.997081 |  |
| STAT5A | K562 |  | 3.488723 | 1.399948 |  |
| TAF1 | Gm12878 |  |  | 8.279398 |  |
| TAF1 | K562 |  |  | 9.316957 |  |
| TAF7 | K562 |  |  | 5.370406 |  |
| TAL1 | K562 |  |  | 2.184837 |  |
| TBL1XR1 | K562 |  | 5.677345 | 3.181479 |  |
| TCF12 | Gm12878 |  | 7.657051 | 4.972183 | 5.183052 |
| TCF3 | Gm12878 |  | 8.015065 | 5.357886 |  |
| TEAD4 | K562 |  | 3.876757 | 1.975382 |  |
| THAP1 | K562 |  | 8.715861 | 8.673127 |  |
| TRIM28 | K562 |  | 3.201460 | 2.129453 |  |
| UBTF | K562 |  | 10.112320 | 11.750788 | 7.112115 |
| USF1 | Gm12878 | 7.276225 |  | 7.753492 |  |
| USF1 | K562 | 5.924492 | 8.185294 | 8.175397 |  |
| USF2 | Gm12878 | 6.325652 |  | 7.796670 |  |
| USF2 | K562 | 5.404733 | 5.701908 | 7.890269 |  |
| YY1 | Gm12878 |  |  | 8.853913 |  |
| YY1 | K562 |  |  | 9.869987 |  |
| ZBTB33 | Gm12878 |  |  | 5.170867 |  |
| ZBTB33 | K562 | 5.630235 | 8.807290 | 5.942027 |  |
| ZBTB7A | K562 | 7.356040 | 12.470241 | 14.281782 | 9.150920 |
| ZEB1 | Gm12878 | 4.603663 |  | 8.050429 |  |
| ZNF143 | Gm12878 |  |  | 9.192158 |  |
| ZNF143 | K562 |  |  | 7.489105 | 9.983317 |
| ZNF263 | K562 |  | 10.278232 | 9.203515 | 9.407070 |

